# Structure and Function of an in vivo Assembled Type III-A CRISPR-Cas Complex Reveal Critical Roles of Dynamics in Activity Control

**DOI:** 10.1101/2021.01.27.428455

**Authors:** Sagar Sridhara, Jay Rai, Charlisa Whyms, Walter Woodside, Michael P Terns, Hong Li

## Abstract

The small RNA-mediated immunity in bacteria depends on foreign RNA-activated and self RNA-inhibited enzymatic activities. The multi-subunit Type III-A CRISPR-Cas effector complex (Csm) exemplifies this principle, but its molecular basis for regulation remains unexplained. Recognition of the foreign RNA, or cognate target RNA (CTR), triggers its single-stranded deoxyribonuclease (DNase) and cyclic oligoadenylate (cOA) synthesis activities. The same activities remain dormant in the presence of the self-RNA, or noncognate target RNA (NTR) that differs from CTR only in its 3’-protospacer flanking sequence. Here we captured four structures of *in vivo* assembled *Lactococcus lactis* Csm (LlCsm) by electron cryomicroscopy representing both the active and the inactive states. Surprisingly, in absence of bound RNA, LlCsm largely forms a minimal assembly lacking the Csm2 subunit with a stably bound catalytic subunit Csm1. Comparison of the minimal LlCsm structure and activities, both in vitro and in vivo, with those of fully assembled LlCsm reveals a molecular mechanism responsible for the viral RNA-activated and self RNA-inhibited activity of Csm1 through protein dynamics.

**Graphic Art Summary:** 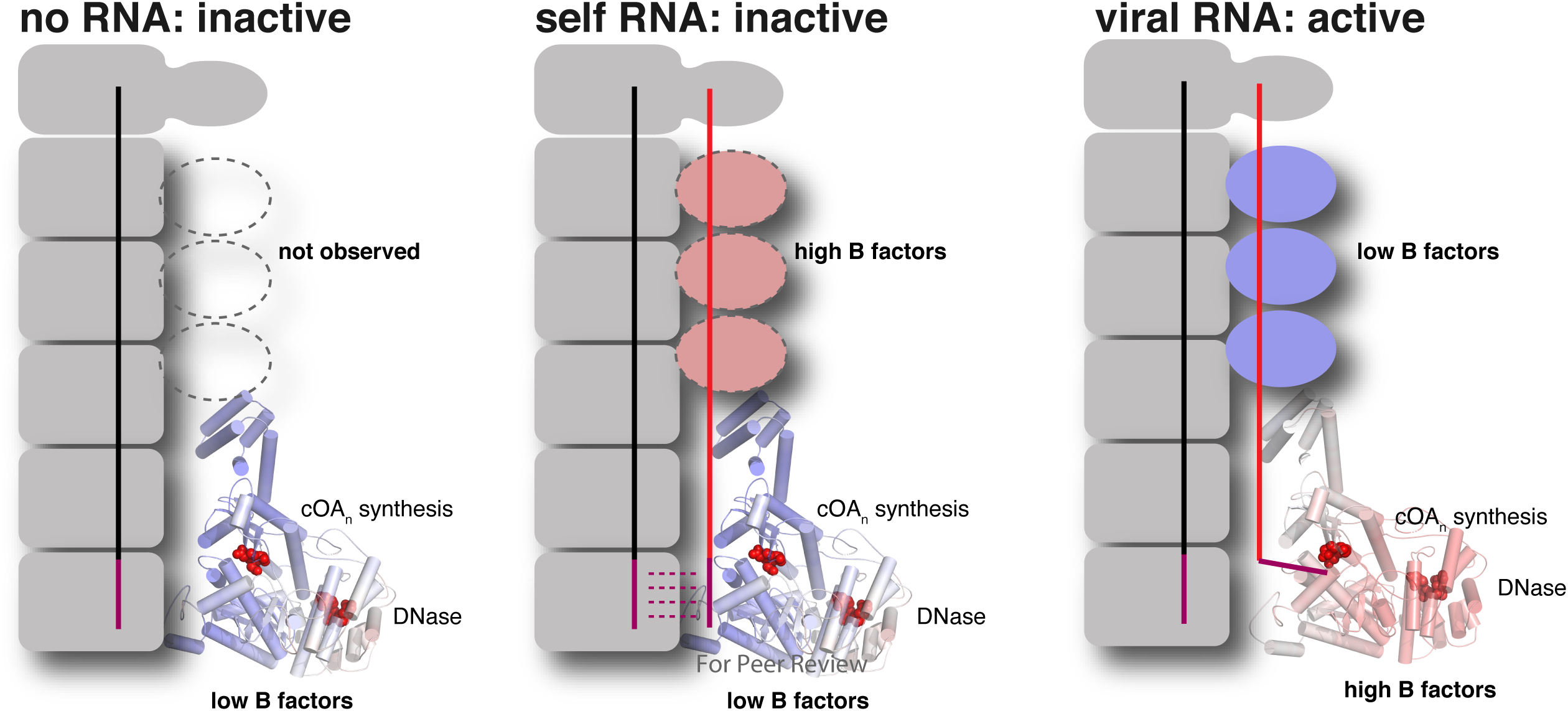

## Introduction

Small RNAs play a wide range of functional roles in microbiota, from transcription termination, stress response, metabolic regulation to immunity through interacting with protein, nucleic acid, and small molecule partners (1–3). Among these, the emerging CRISPR (<u>C</u>lustered <u>R</u>egularly Interspaced <u>S</u>hort <u>P</u>alindromic <u>R</u>epeats) RNA-based immunity affords the most complex processes of RNA regulation. CRISPR is a characteristic genetic feature of bacteria and archaea required to mount immunity against invading nucleic acids originating from viral infections or invasion of plasmids or other mobile genetic elements (4–8). CRISPR partners with CRISPR-associated (Cas) proteins to interfere against the invading nucleic acids. Currently known CRISPR-Cas systems are remarkably broad both in composition and interference mechanisms. Understanding the molecular basis of CRISPR-Cas systems not only unravels the most fundamental RNA-mediated biochemical mechanisms underlying the microbiological warfare but also leads to novel biotechnological applications (9, 10).

Among the known types of CRISPR-Cas systems, the Type III is unique in that it elicits multifaceted immune responses. Type III systems are triggered by actively transcribing viral messenger RNA (mRNA) that bears sequence complementarity to the CRISPR RNA (crRNA) carried by Type III effectors. Upon activation, Type III systems directly degrade invader mRNA and its encoding DNA, and more surprisingly, synthesize cyclic oligoadenylates (cOAs) that generate secondary antiviral responses (11). The molecular mechanism of Type III systems is, therefore, arguably the most complex among all types of known CRISPR-Cas systems. Based in part on features of the largest and the signature subunit: Cas10, the Type III systems are further divided into four subtypes: Type III-A/D (Csm), Type III-B/C (Cmr), Type III-E and Type III-F (12), among which the Csm CRISPR-Cas effector complex has been studied most extensively by both biochemical and structural methods (13–16).

Csm is comprised of five proteins: Cas10, or Csm1, Csm2-Csm5 and a crRNA that form a “seahorse”-shaped assemblage. The single-stranded DNase (ssDNase) and the cOA synthesis activities reside within the Csm1 subunit (17, 18) and the RNase activity resides within Csm3 (19–24). While the Csm3-mediated RNase activity requires the mere crRNA-target RNA complementarity (25), the Csm1-mediated DNase and cOA synthesis are allosterically regulated (26, 27). Binding of a cognate target RNA (CTR) that complements the crRNA guide region but not the 8-nucleotide (nt) repeat-derived 5’-crRNA tag sequence (alternatively referred to as a 5’-handle), induces non-specific DNase and the cOA synthesis activities (16,18,28,29). In contrast, a target RNA that complements both the guide region and the 5’-tag of the crRNA (henceforth referred to as the non-cognate target RNA, or NTR) is unable to switch on the Csm1-mediated activities (26). In other words, NTR represents “self” RNA and thus prevents Csm-mediated autoimmunity. Interestingly, cleavage of CTR by Csm3 causes cessation of the Csm1-mediated activities (26), thereby providing a temporal regulation of DNase activity that avoids the potential cleavage of the host genome (18, 30). Furthermore, while isolated Csm1 is a constitutively active DNase (31, 32), the Csm assembly (apo effector) possesses no such activity, indicating that both Csm assembly as well as target RNA regulate the enzymatic activities of Csm1.

The extraordinary control of Csm1 activities remains partially understood. Previous systematic structural studies have provided detailed three-dimensional views of Csm complexes from bacterial and archaeal species in various functional states that include the apo, the NTR-bound, the CTR-bound, and the NTR/CTR-bound with nucleotides (13,14,16,33–35). In addition, high resolution crystal structures of individual subunits or subcomplexes from a number of species with or without bound nucleotides are known (32,36,37). These structural data and the associated biochemical analyses have created an exceptional opportunity to begin to resolve the molecular mechanism of the Csm. The observed structural transitions from the apo to the NTR- or the CTR-bound state provide reasonable explanations for the activity and regulation of the RNase, but less satisfactorily for the cOA synthesis and the DNase activities. As a result, structural dynamics was proposed to be a method of regulation (13,14,36), which has since found experimental support (26). The single-molecule fluorescence study showed that in the presence of CTR and NTR, respectively, Csm1 exhibits different conformational dynamics (26). However, the underlying mechanism of dynamic regulation remains largely uncharacterized.

The single-particle cryoEM method is a powerful method for providing atomic structures and dynamic information of the imaged specimens (38–40). Although one must take into the consideration of the specimen preparation process and radiation damage, cryoEM captures the entire inventory of states from a single enzymatic complex that can represent a snapshot of its conformations in solution. Conformational fluctuation is reflected as structure heterogeneity and results in weak electron potential density and thus high atomic displacement parameters in the refined models. Statistical methods have been developed for further elucidating the nature of structural heterogeneity if all or part of the protein unit can be modeled by rigid-body motions (39). These structural parameters can be used to identify the site and the nature of dynamic motions.

The *Lactococcus lactis* gram positive bacterium is among the most extensively used microorganisms in dairy industry and possesses a range of medical and biotechnological applications (41, 42). Although *L. lactis* was thought to rely only on various non-CRISPR-mediated defense mechanisms (43), a novel conjugative plasmid-encoded type III-A CRISPR-Cas system (Figure 1A) was identified (44). The plasmid-encoded *L. lactis* Csm (LlCsm) system was subsequently shown to confer a similar transcription-dependent plasmid DNA interference activity to those by its orthologs from *S. thermophilus*, *S. epidermidis* and *T. thermophilus* (45, 46). We reconstituted LlCsm complexes *in vivo* and demonstrated its enzymatic activities. We report four cryo-EM structures of the Csm2-free apo, the NTR-, and the CTR-bound full LlCsm complexes that collectively represent the active and the inactive states. Structural analysis identified an inversely correlated structural dynamics between the catalytic subunit Csm1 and the target RNA-binding Csm2 in a function-dependent manner. Mutagenesis, fluorescence and plasmid interference assay were performed that support an RNA-regulated dynamic model for regulation.

**Figure 1.**
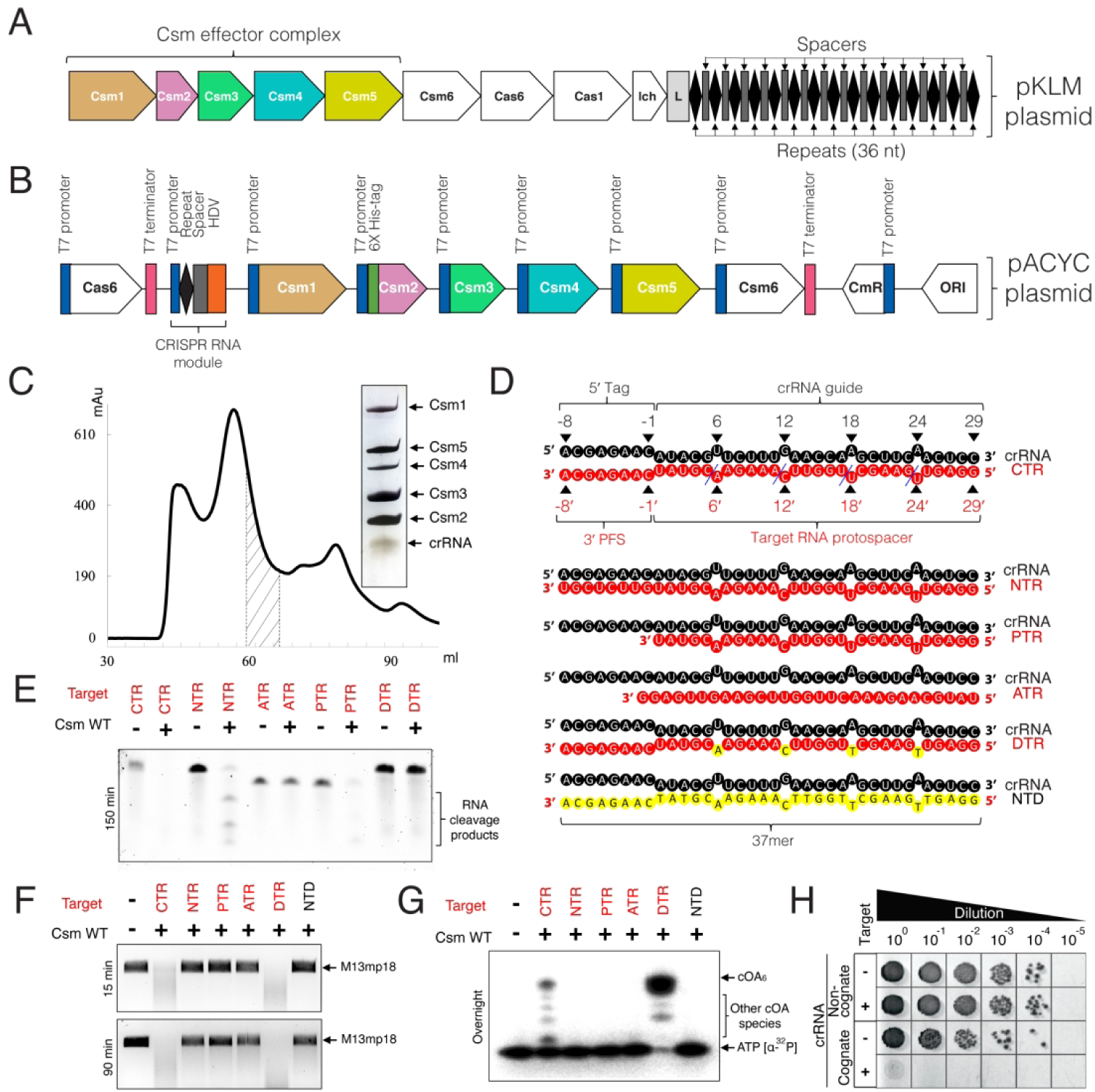
Purification and activity of *Lactococcus lactis* type III-A (Csm) CRISPR-Cas system. (A) Genome organization of Type III-A CRISPR-Cas system in pKLM plasmid of *Lactococcus lactis*. The locus harbors tandem copies of 36 nt repeats (diamonds) and spacers (rectangles) of varying length. (B) Schematic representation of pACYC expression plasmid harboring *L. lactis* CRISPR-Cas genes Csm1-6, Cas6 and CRISPR-RNA locus in an HDV-ribozyme construct. (C) Size-exclusion chromatography and Silver stain profile of purified LlCsm RNP. The fractions pooled are indicated by shaded region. (D) Schematic representation of crRNA and various crRNA-target RNA duplexes used in the study (crRNA: CRISPR RNA, CTR: Cognate target RNA, NTR: Non-cognate target RNA, PTR: Protospacer target RNA, ATR: Anti-sense target RNA, DTR: Deoxy-substituted target RNA, NTD: Non-target DNA). The blue dashed lines indicate target RNA cleavage sites. (E) *In vitro* RNA cleavage assay by WT LlCsm complex. The RNase reactions were incubated for 150 min. (F) *In vitro* DNA cleavage assay by WT LlCsm complex. M13mp18 was used as substrate and the reaction products were analyzed at timepoints 15 min and 90 min. (G) *In vitro* cOA synthesis assay by WT LlCsm complex. Radiolabeled α-^32^P-ATP was used as a substrate and the reactions were incubated overnight. (H) *In vivo* plasmid interference assays with the wild type LlCsm harboring a non-cognate crRNA or the wild type LlCsm harboring a cognate crRNA.

## Material and Methods

### Protein expression and purification

The pACYC *Lactococcus lactis* module plasmid encoding proteins Cas6, Csm1-6 and CRISPR locus was as described previously (Figure 1B, Tables S3 and S4) (46). The crRNA locus in the plasmid was suitably modified to contain repeat-spacer-HDV ribozyme sequence (Figure 1), so that Cas6 and HDV ribozyme would perform 5’- and 3’-crRNA processing, respectively. The *csm2* gene encoded an N-terminal His_6_-tag in all variants of LlCsm. The ΔCsm2-LlCsm was produced by Gibson assembly such that csm3 gene encoded an N-terminal His_6_-tag. The mutations were introduced by Q5 mutagenesis (New England Biolabs) and verified using sequencing primers (Eurofins Genomics). All variants of LlCsm ribonucleoprotein complex were produced in *Escherichia coli* NICO strain using 0.3 mM isopropyl β-D-1-thiogalactopyranoside (IPTG) for induction of protein expression. The RNP was purified using nickel-affinity chromatography followed by size-exclusion chromatography in 20 mM HEPES pH 7.5, 200 mM NaCl, 5 mM MgCl_2_, 14 mM 2-mercaptoethanol. The fractions of the final gel filtration step containing all five Csm proteins associated with crRNA were pooled, concentrated, aliquoted and flash frozen using liquid nitrogen before storage in - 80 °C (Figure 1C, Figure S1B).

The His_6_-tagged LlCsm6 (Figure 7A) was separately produced for activity assays in *Escherichia coli*. LlCsm6 was purified using nickel-affinity chromatography followed by size-exclusion chromatography in the same buffer used for LlCsm. The fractions containing homogenous LlCsm6 from the final gel filtration step were pooled, concentrated, aliquoted and flash-frozen using liquid nitrogen before storage in −80 °C.

### In vitro assembly of RNP-target RNA complex

The 45mer NTR/CTR target RNA (Figure S1A, Table S2) with 29 bases crRNA complementarity was ordered from Integrated DNA Technologies (IDT). The purified LlCsm (D14N)_1_(D30A)_3_ effector complex was mixed with NTR/CTR at 1:2 molar ratio, incubated at 37 °C for 60 min and resolved using analytical Superdex S200 column (GE Healthcare) in a buffer containing 20 mM HEPES pH 7.5, 200 mM NaCl, 5 mM MgCl_2_, 14 mM 2-mercaptoethanol (Figure S1C). The peak fractions containing target-bound LlCsm complex (Figure S1D) was used to prepare cryo-EM grids. The preparation of apo-Csm complex followed the same buffer and pre-incubation conditions with the exception of no target RNA was added (Figure S1D).

### Cryo-EM sample preparation

The LlCsm RNP and RNP-target RNA samples at approximately 0.5 mg/ml were separately applied in 4 μL volume onto glow-discharged UltrAuFoil 300 mesh R1.2/1.3 grids (Quantifoil), blotted for 3 s at 88% humidity and flash-frozen in liquid ethane using FEI Vitrobot Mark IV. The grids were stored in liquid nitrogen before being used for imaging.

### EM data collection, processing, and 3D reconstruction

The ice-embedded LlCsm complex samples were collected on FEI Titan Krios electron microscope equipped with Gatan Bioquantum K3 direct electron detector (ThermoFisher Scientific) with the Leginon software for automatic data acquisition (47) in a counting mode. Both motion correction and contrast transfer function (CTF) estimation were performed in RELION 3.0 (48) that provides the UCSF MotionCor2 (49) and Gctf wrapper (50). Particles were auto picked using the LoG-based auto-picking algorithm implemented in RELION-3 (48). The stack was created and imported into cryoSPARC (51) for 2D classification in order to eliminate bad particles. The resolution was estimated using the gold-standard Fourier Shell Correlation (FSC) plot at the value of 0.143. Local resolution was estimated using Resmap (52). All images were collected at 81,000 x magnification with 1.074 Å/pixel sampling rate at the specimen level and −1.3 to −2.8 microns defocus. A 60.07 e^−^/Å^2^-61.51 e^−^/Å^2^ dose was applied over 70-74 frames at a total exposure time of 3.09 s - 4.35 s.

A total of 5,179 images were collected from the CTR-bound complex in a movie mode (Figure S3). Images showing bad ice, astigmatism, drift, and poor sample quality were rejected resulting in 2,543 images for further processing and particle picking, which resulted in a total of 2,157,655 particles. Several rounds of 2D classification led to 613,110 particles with good quality. RELION-3 (48) was used to classify the particles, which led to further reduction of particles to 229,062 based on high resolution features. The classes with similar features were combined and auto-refined using a custom 3D mask that resulted in an overall resolution of 3.57 Å. Multiple rounds of ctf refinement were performed on RELION 3.1 (53), which improved resolution to 3.19 Å. To further resolve heterogeneity near the top of the particles, classification without alignment was performed using a mask around the Csm5 subunit, resulting in three major classes. The largest class shows no bound target RNA nor Csm2 subunit densities and therefore, no further refinement was carried out. Two other classes show strong densities of the bound target RNA and are refined to two structures with different stoichiometry (54,783 particles, CTR-43 and 62,413 particles, CTR-32). The CTR-43 class and CTR-32 class were independently further refined in RELION 3.1 (53) to a resolution of 3.27 Å and 3.35 Å respectively. The resolution of class CTR-43 was improved to 3.07 Å with further autorefinement with cisTEM (54).

A total of 4,382 images were collected from the apo complex in a movie mode (Figure S4) that resulted in 2,336 after rejection of bad images. A total of 3,636,087 particles were auto picked and 2D classification was performed to remove the bad particles, which led to 1,290,536 particles. Classification with RELION-3 (48) further reduced the particles to 436,641 based on high resolution features. The classes with similar features were combined and refined by iterative auto-refinement using a custom 3D mask and followed by multiple rounds of CTF refinement that resulted in an overall resolution of 2.98 Å. Refinement with cisTEM (54) of the same stack improved the resolution to 2.97 Å.

A total of 3,922 images were collected from the NTR-bound complex in a movie mode (Figure S5) that resulted in 1,892 after rejection of bad images. A total of 2,865,726 particles were auto picked and 2D classified to remove the bad particles, which led to 1,036,382 particles. 3D Classification with cryoSPARC (51) further reduced the particles to 282,878 based on high resolution features. The classes with similar features were combined for further classification that separated the classes with and without target RNA. The classes with target RNA were combined and refined by iterative auto-refinement in RELION 3.0 (39) resulted in an overall resolution of 3.57Å. Furthermore, like CTR refinement, classification without alignment was performed using a mask around the Csm5 subunit, to solve the heterogeneity resulting in three major classes. Two classes show no density of target RNA, and therefore no further refinement was carried out. The remaining class (39,220 particles) shows the target RNA density with four copies of Csm3 and three copies of Csm2. This class was further refined multiple rounds with CTF refinement in RELION 3.1 (53) that resulted in 3.48 Å resolution.

### Multibody Refinement

Multi-body refinement was carried out for the Apo, CTR-43, and NTR-bound complex, respectively, using RELION-3 (48) by defining three bodies (HD domain, Csm1 without HD, and the rest of Csm1), which revealed a substantial movement of Csm1 with respective to the rest of the complex. In all cases, the first three eigenvectors represent more than 50% of the particle motion and are shown in Figure 6. Densities were segmented by the watershed segmentation feature available in UCSF Chimera (55).

### Model Building and Refinement

The highest resolution map of the apo complex was used to build models for Csm1, Csm3, Csm4 and Csm5 and the crRNA starting from their homology-build models and manually adjusted in COOT (56). The map of the CTR_43 complex was used to build Csm2 and the CTR and that of the NTR_43 was used to build NTR. Real-space refinement was carried out for all models with PHENIX (57). For all complexes, the RELION-3 maps were first used in model building and refinement. During early stage of the refinement, B factors for all atoms were set arbitrarily to 20.0 Å^2^. Iterative rounds of real-space refinement and manual building in COOT led to models with overall correlation coefficients to be greater than 0.65. In the final stages of the refinement, the atomic displacement parameters (B-factors) were refined with iterative rounds of positional refinement, leading to models with overall correlation coefficients to be greater than 0.8 with excellent stereochemistry parameters. These models were used to compute Q-scores with the MapQ plugin (58) in UCSF Chimera (55). The final models were refined against the highest resolution electron potential density either with cisTEM or RELION-3.1 (Table S2).

### In vitro target RNA cleavage assays

The RNA cleavage assays were performed in a cleavage buffer containing 33 mM Tris acetate pH 7.6/32 °C, 66 mM potassium acetate, 10 mM MnCl_2_. The reactions were performed at 37 °C/30 to 150 min as indicated and contained 100 nM LlCsm complex and 500 nM target RNA (Table S3). The reactions were quenched using 2x formamide dye (95% formamide, 0.025% SDS, 0.025% bromophenol blue, 0.025% xylene cyanol FF, 0.5 mM EDTA). The reaction products were heated at 70 °C/3 min and separated by 7 M Urea, 15% polyacrylamide-gel electrophoresis (PAGE) gels in 1x Tris/Borate/EDTA (TBE) running buffer and were visualized by staining with SYBR Gold II (Invitrogen) stain.

### In vitro DNA cleavage assays

The DNA cleavage assays were performed as previously described (17). Briefly, 2 nM M13mp18 circular ssDNA (New England Biolabs) (Table S3) was treated with 200 nM LlCsm complex, 200 nM target RNA at 37 °C/10 to 90 min as indicated in a cleavage buffer containing 33 mM Tris acetate pH 7.6/32 °C, 66 mM potassium acetate, 10 mM MnCl_2_. The reactions were quenched using 1x purple gel loading dye (New England Biolabs). The reaction products were heated at 95 °C/5 min and separated on 1% agarose gel in Tris/Acetic acid/EDTA (TAE) running buffer and were visualized by staining with Ethidium Bromide.

### In vitro cOA synthesis assays

The cOA synthesis assays were performed as previously described (59). Briefly, a mixture containing 160 μCi α-^32^P-adenosine triphosphate (PerkinElmer) and 500 μM ATP was incubated with 100 nM LlCsm complex, 200 nM nM target RNA at 37 °C/overnight in a cOA synthesis buffer containing 33 mM Tris acetate pH 7.6/32 °C, 66 mM potassium acetate, 10 mM MgCl_2_. The reaction products were heat denatured at 95 °C/10 min and centrifuged at high speed. The supernatant was mixed with formamide dye, resolved by 8 M Urea, 24% PAGE gels in 1x Tris/Borate/EDTA (TBE) running buffer at 80 V/240 min. The gels were carefully sandwiched between non-porous cellophane sheet and a porous gel drying sheet and dried for 120 min, developed for 30 min using Phosphor Screen (GE Healthcare Life Science) and visualized using the Typhoon Gel Imaging System (GE Healthcare Life Science).

### Fluorescence reporter assays

The fluorescent DNA reporter assay was performed using a DNA probe (IDT) covalently linked to 5’-Alexa Fluor 594 (NHS Ester) fluorescent dye and 3’-Iowa Black RQ quencher (Table S3). The kinetic process was monitored by the addition of the target RNA (500 nM) to a mixture containing 250 nM LlCsm effector complex and 1000 nM substrate DNA probe in a buffer containing 33 mM Tris acetate pH 7.6/32 °C, 66 mM potassium acetate, 10 mM MnCl_2_ at 37 °C. The fluorescence was measured on Spectramax ID5 multi-mode microplate reader (Molecular Devices) using 570 nm/630 nm excitation/emission wavelengths at 60 s intervals. The reactions were performed in triplicates and averaged for the final plots. The fluorescent RNA reporter assay was performed using an RNA probe (IDT) covalently linked to 5’-6-FAM (Fluorescein) fluorescent dye and 3’-Iowa Black FQ quencher (Table S3). The reactions was initiated by the addition of 500 nM target RNA to a mixture containing 1000 nM RNA probe, 250 nM LlCsm effector complex, 1 nM LlCsm6, 1000 nM substrate RNA probe in a buffer containing 33 mM Tris acetate pH 7.6/32 °C, 66 mM potassium acetate, 10 mM MgCl_2_, 10 mM MnCl_2_, 0.5 mM ATP at 37 °C and was measured on Spectramax ID5 multi-mode microplate reader (Molecular Devices) using 480 nm/530 nm excitation/emission wavelengths at 60 s intervals. The reactions were performed in triplicates and averaged for the final plots.

### In vivo plasmid interference assays

Plasmid interference assays were carried out as previously described (46). Chemically competent BL21-AI *E. coli* were transformed with pCsm variants and transformants were selected on Miller’s LB broth (Invitrogen) agar supplemented with 34 μg/ml chloramphenicol. Single colonies were cultured in super optimal broth medium (SOB) (BD Difco) and made electrocompetent through successive washes with 10% glycerol. In triplicate, 100 ng of pTrcHis plasmid (with or without transcribed complementary target sequences) were added to 50 μl of competent cells in a 0.2 cm-gap Gene Pulser® electroporation cuvette (BioRad) on ice. Cuvettes were transferred to a Gene Pulser II (BioRad) and pulsed with the following settings: 25 μF capacitance, 2.5 kV, and 200 ohms. Immediately following transformation, 950 μl of super optimal broth with catabolite repression (SOC) (SOB + 20 mM glucose) was used to recover pulsed cells from the cuvette and moved to 1.5 ml microcentrifuge tubes. The tubes were shaken at 200 rpm at 37 °C for 60 minutes. Serial 10-fold dilutions were made to 10-5 and spot plated onto LB agar containing 100 μg/mL ampicillin and 34 μg/mL chloramphenicol to select for pTrcHis and pCsm, respectively. Plates were imaged after overnight incubation at 37 °C.

## Results

### Reconstitution of Functional LlCsm In vivo

Csm complex contains five proteins and one guide crRNA that have been previously observed to assemble into ribonucleoprotein (RNPs) particles of variable stoichiometries (13, 21). In order to produce LlCsm RNPs that represent those present in *L. lactis*, we employed a multi-protein-RNA co-expression system: a previously constructed single pACYC-module plasmid (46), that encodes Cas6, Csm1-Csm5, a precursor crRNA (pre-crRNA) followed by the hepatitis delta virus (HDV) self-cleaving ribozyme, and Csm6 locus (Figure 1B). Co-expression and concerted processing of the repeats by Cas6 endoribonuclease and of the spacer by the HDV ribozyme gave rise to a mature crRNA of 37-nt bound to LlCsm complex (Figure 1C & Figure S1). The N-terminally His-tagged Csm2 subunit enabled the isolation of the entire LlCsm complex via metal-affinity and size exclusion chromatography. The final homogenous LlCsm complex was verified to contain Csm1-5 and the 37-nt crRNA (Figure 1C & Figure S1) and was used in subsequent activity assays and structural studies. As previously observed (60), Csm6 is not a stable component of the Csm complex (Figure 1C).

Similar to what was observed for other homologous Csm complexes, the *in vivo* assembled LlCsm possesses Csm3-mediated target RNA cleavage (Figure 1E), the Csm1 HD domain-mediated ssDNA cleavage (Figure 1F), the Csm1 Palm2 domain-GGDD motif mediated cOA synthesis (Figure 1G) and plasmid interference activities (Figure 1H). Specifically, LlCsm cleaves any target RNA that is complementary to the guide region of the crRNA (Figure 1E). The LlCsm ssDNase is activated only in the presence of CTR and remained dormant in the presence of NTR (Figure 1F). Other target RNAs such as a non-complementary, anti-sense target RNA (ATR) or a protospacer target RNA (PTR) that lacks the 8-nt PFS, failed to elicit ssDNA cleavage (Figure 1F) or cOA synthesis activity (Figure 1G). Noteworthy, the deoxy target RNA (DTR) whose cleavage site 2’-hydroxyl groups are replaced by hydrogen, induced greater extent of ssDNA cleavage (Figure 1F) and cOA synthesis (Figure 1G), consistent with previous observations that cleavage of CTR reduces ssDNA cleavage (18, 30).

Consistent with the *in vitro* DNase and cOA synthesis activities, bacterial cells transformed with the plasmid encoding the cognate LlCsm RNP and LlCsm6 resulted in plasmid interference and loss of cell growth when the transcribed target sequence was present but not in its absence (Figure 1H, Table S1, Figure S2). Mutation in the HD domain of LlCsm1 or that of the RNase center of LlCsm3 (Asp30) did not impact while mutation of the Palm2 domain was deleterious to the interference activity (Table S1, Figure S2), suggesting a more important role of cOA synthesis than either the DNase or the RNase in interfering the target plasmid in cells as observed previously (60). The critical importance of the cOA_n_-activated Csm6 RNase in immunity is consistent with the previous observation that activated Csm does not directly cleave the unwound DNA bubble in the transcription elongation complex (16).

### Distinct LlCsm Structural Assemblies

We employed cryoEM to determine the structures of the LlCsm complexes alone or bound with the NTR or CTR forms of the target RNA. The LlCsm complexes harboring Csm1 Asp14Asn and Csm3 Asp30Ala mutations (13, 14) were used in cryo-EM studies in order to prevent degradation of the bound nucleic acids. The apo, the NTR-bound and the CTR-bound complexes were prepared similarly prior to making cryo-grids. The co-purified LlCsm complex was pre-incubated with or without CTR or NTR in molar excess (RNP : Target RNA = 1:2) followed by fractionation on a size exclusion column (Figure S1). The peak fractions of each run containing the complex were suitably diluted before plunging to achieve homogenous particle distribution. Data collection and single particle reconstruction resulted in four main structures: the apo structure (2.9 Å overall), the NTR-bound structure (3.4 Å overall) and the two CTR-bound structures with different subunit stoichiometry (3.0 Å and 3.3 Å overall), wherein the two CTR-bound structures were distinct classes of the CTR-bound specimens (Materials and Methods).

The electron potential densities for Csm3, Csm4, crRNA and the target RNA, if bound, have excellent quality in all structures to allow tracing and placement (Figures S3, S4, S5 & S6). The full-length Csm1 could be reliably traced in both the apo or the NTR structure while focused refinement with a Csm1 mask in the CTR structure allowed its placement (Figure S6). A large majority of Csm5 was also traced in all three structures. The clear but weaker electron potential densities for both Csm1 and Csm5 in the CTR-bound structures is indicative of their dynamic motion in these complexes (see more below). All final structures were refined to satisfactory correlation coefficients and geometry (Table S2).

Two distinct CTR-bound structures were resolved from two major classes that have 1_1_2_3_3_4_4_1_5_1_crRNA_1_Target_1_ (CTR-43) and 1_1_2_2_3_3_4_1_5_1_crRNA_1_Target_1_ (CTR-32) stoichiometry, respectively, where each number corresponds to a numbered Csm protein (Figure 2, Figure S3). We describe overall features of LlCsm on the higher-resolution CTR-43 complex. The portion of the CTR-32 complex that is common with CTR-43 does not differ from CTR-43 in structure (Figure 2B). LlCsm is essentially comprised of two helical ridges capped at the foot by the characteristic subunit Csm1. One ridge is made of RAMP (<u>R</u>epeat <u>A</u>ssociated <u>M</u>ysterious <u>R</u>epeat) protein or domains (R ridge) and the other is made of helical proteins (H ridge). Four Csm3 subunits, the RAMP domain of Csm4 (at the foot) and that of Csm5 (at the head) comprise the R ridge whereas three Csm2 subunits, the helical domain (domain D4) of Csm1 (at the foot) and that of Csm5 (at the head) comprise the H ridge (Figure 2). The interface between each pair of RAMP subunits/domain buries a large solvent accessible area (∼1266 Å^2^) with nearly equal numbers of hydrogen bonds (8–11) and salt bridges (7–9) and significant non-polar interactions (Figure 2). The only exception is the interface between the RAMP domain of Csm5 and Csm3.4 that has a smaller solvent accessible area (∼994 Å^2^) and only one salt bridge, reflecting a weaker interaction between Csm5 and the complex. The H ridge has interfaces of ∼half the buried solvent accessible area and less number of salt bridges than those of the R ridge (Figure 2). At the foot of the two ridges, Csm4 interacts with Csm1 extensively with 1261 Å^2^ buried solvent accessible area, 14 hydrogen bonds, and 4 salt bridges (Figure 2). The crRNA traverses along and buries a large interface with the R ridge (Figure 2). The CTR, in contrast, traverses along and buries a relatively small interface with the H ridge (Figure 2). The smallest buried surface area is found between pairs of Csm3 and Csm2 subunits (∼420 Å). These interfaces have 3-6 hydrogen bonds but no salt bridges. As interaction strength is correlated with the amount of buried solvent accessible area (61), the above analysis indicates that the R ridge and its interaction with crRNA is thus more stable than the H ridge.

**Figure 2.**
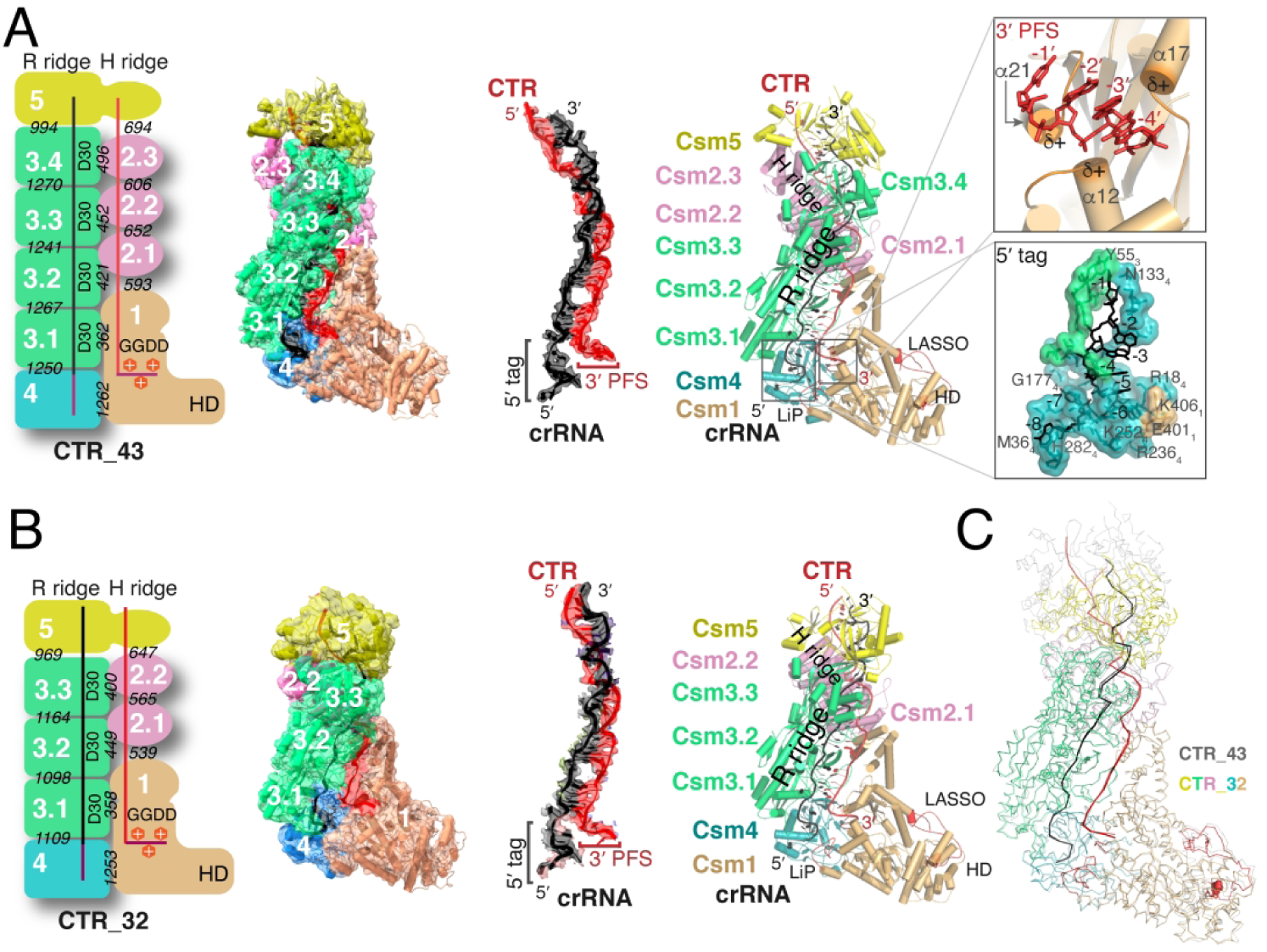
Structures of Cognate target RNA (CTR)-bound LlCsm complex. (A) Schematic representation, electron potential density, and cartoon representation of CTR-43 with stoichiometry 1_1_2_3_3_4_4_1_5_1_crRNA_1_Target_1_. Numbers in while indicate names of the subunit where N or N.m represent CsmN (or the mth of CsmN). Italicized numbers in black indicate the buried solvent accessible surface of the corresponding subunits in Å^2^. The crRNA and CTR are colored black and red respectively. The relative positions of crRNA 5’-tag and the CTR 3’-PFS are highlighted. The boxed image at the top displays three alpha helices of LLCsm1: α12, α17 and α21 that orient the positively charged N-terminus towards the 3’-PFS of CTR, thus forming a favorable protein-RNA interaction. The boxed image towards the bottom displays the Csm3-Csm4 pocket where the 5’-tag of crRNA traverses. (B) Schematic representation and overall structure of CTR-32 with stoichiometry 1_1_2_2_3_3_4_1_5_1_R_1_T_1_. The crRNA and CTR are colored black and red respectively. The relative positions of crRNA 5’-tag and the CTR 3’-PFS are highlighted. (C) Ribbon representation of CTR-43 and CTR-32 with their Csm3 subunits superimposed highlighting structural differences in other subunits.

Despite Csm2 being the His-tagged subunit, it was surprisingly missing from the apo structure except for some weak density near Csm1, resulting in a 1_1_2_0_3_4_4_1_5_1_crRNA_1_Target_0_ stoichiometry (Figure 3, Figure S4). Since all Csm subunits were present immediately before making the cryoEM grids (Figure S1), the reason for the lack of Csm2 in the apo complex is likely due to its dissociation from the complex during plunge freezing. Nearly all particles that went into reconstruction of the apo structure contain no Csm2 (Figure S4). On the other hand, the NTR-bound LlCsm complex has excellent density for all subunits despite its slightly lower overall resolution, resulting in the 1_1_2_3_3_4_4_1_5_1_crRNA_1_Target_1_ stoichiometry (Figure 4 & Figure S5). Notably, both the CTR-bound and the NTR-bound particles contain a large class without a target RNA that also lacks the LlCsm2 subunits (Figures S3 & S5), further supporting the lack of or weak Csm2 association with the apo complex observed in the ice-embedded samples.

**Figure 3.**
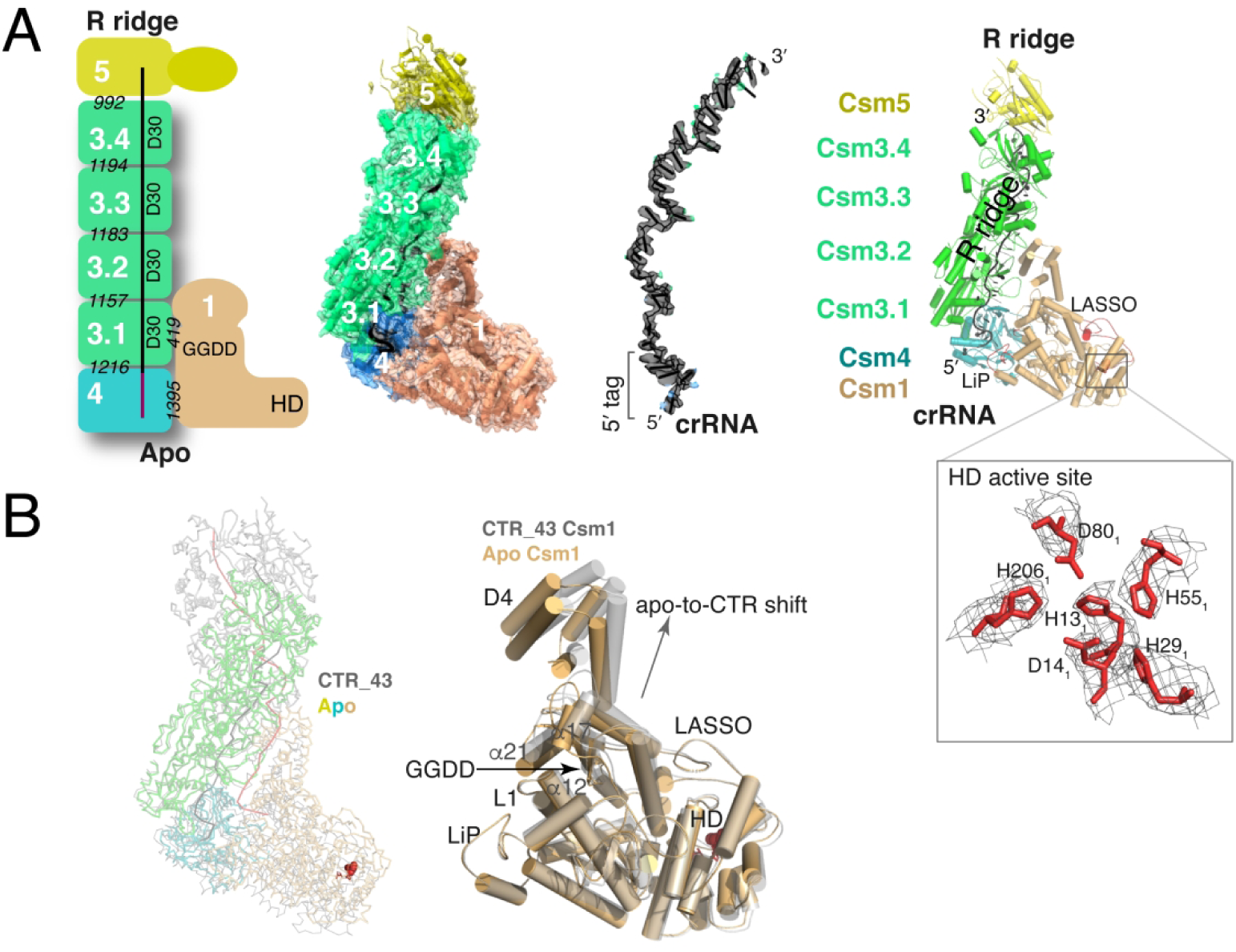
Structure of apo LlCsm complex lacking a bound target RNA. (A) Schematic representation, electron potential density and cartoon representation of the LlCsm apo complex with stoichiometry 1_1_2_0_3_4_4_1_5_1_crRNA_1_Target_0_. Numbers in while indicate names of the subunit where N or N.m represent CsmN (or the mth CsmN). Italisized numbers in black indicate the buried solvent accessible surface of the corresponding subunits in Å^2^. The crRNA is shown in black. The boxed image shows close-up view of catalytic HD residues with electron potential density. The putative catalytic residues His13, Asp14, Asp80 and His206 form a permuted HD motif and are strongly conserved. (B) Ribbon representation of apo LlCsm complex (colored) overlaid with LlCsm CTR-43 complex (gray) using their Csm3 subunits. The HD residues are highlighted in red. The cartoon representation depicts upward shift of Csm1 D4 domain upon binding to CTR wherein alpha helices α12, α17 and α21 follow along. Note the minimal shifts among LASSO, LiP and L1 loops.

**Figure 4.**
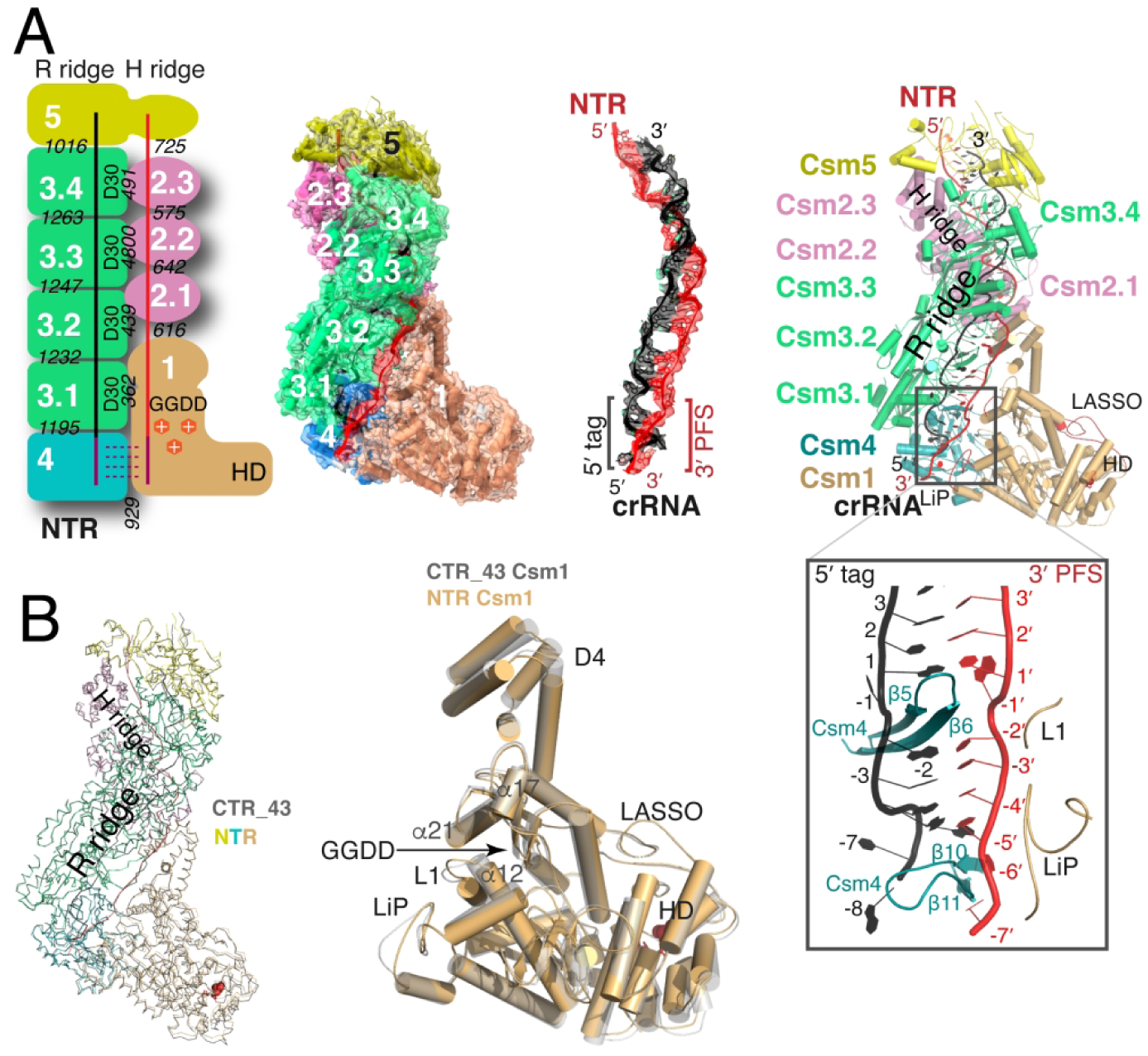
Structure of the noncognate target RNA (NTR)-bound LlCsm complex. (A) Schematic representation and electron potential density of NTR complex with stoichiometry 1_1_2_3_3_4_4_1_5_1_crRNA_1_Target_1_. Numbers in while indicate names of the subunit where N or N.m represent CsmN (or the mth CsmN). Italicized numbers in black indicate the buried solvent accessible surface of the corresponding subunits in Å^2^. The crRNA and NTR are shown in black and red, respectively. The relative locations of crRNA 5’-tag and the NTR 3’-PFS are highlighted. The boxed image displays the base paired 5’-tag and 3’-PFS surrounded by Csm4 and Csm1 structural elements. (B). Comparison of CTR-43 and NTR complex in ribbon representations. The Csm3 subunits of the two complexes are superimposed to reveal, if any, differences in other subunits. Csm1 subunits from both complexes are compared in a cartoon representation. Key secondary elements are labeled.

### RNase Centers Link RNase Activity of LlCsm2 to LlCsm1

Previous kinetic analysis showed that both the cOA synthesis and DNase activities dampen with the progression of target RNA cleavage (30). Further, we showed that a non-cleavable target RNA yielded stronger cOA and DNase activities than a cleavable target RNA did (Figures 1F & 1G). To shed light on the mechanistic link between RNA cleavage and Csm1 activities, we analyzed the structure of the RNase centers of LlCsm. Although Csm requires a divalent ion to cleave its target RNA (18, 22), the structural arrangement of the observed target RNA cleavage centers and the fact that the cleavage products of the analogous Type III-B/Cmr complex have 2’,3’-cyclic phosphate termini (62) suggest that it follows RNase A-like cleavage mechanism (63) similar to that of the crRNA processing endoribonuclease Cas6 (64). This reaction mechanism is also consistent with our observed inhibition of cleavage if the 2’-hydroxyl group upstream of scissile phosphate is replaced by hydrogen (Figure 1). Four RNase centers are observed in the CTR-43 complex along the backbone of the bound CTR (Figure 5). The RNase sites are marked by the thumb loop of the Csm3 subunits that harbors the critical Asp30 residue. However, since the Asp30Ala mutant was used to form the cryoEM sample, Asp30 was not observed and the substituted Ala30 is 4-5 Å away from the leaving group oxygen. Despite so, the thumb loop-facilitated flipping of the nucleotide 5’ of the cleavage sites enables a favorable conformation for cleavage. The 2’-nucleophilic oxygen, the scissile phosphate group, and the leaving group oxygen form the classic inline geometry at each of the four sites (Figure 5A). Either a tyrosine (Tyr693 of Csm1, site 1) or an arginine (Arg48 of Csm2, sites 2-4) residue is situated near the 2’-nucleophilic oxygen that could act as the general base to extract the proton (Figure 5B). Consistently, mutation of Tyr693 of Csm1 or Arg48 of Csm2 severely reduced RNA cleavage (Figure 5C). Similarly, mutation of the homologous tyrosine in StCsm1 (Tyr686Ala) (Figure S7) and the arginine in StCsm2 (Arg41Ala) (Figure S8) were also found to diminish or reduce the cleavage activity in the previously studied *Streptococcus thermophilus* Csm (14). A positive charged residue (lysine or arginine) is observed to be placed near the scissile phosphate that could stabilize the negative charge developed on the penta-coordinated phosphate. Consistently, reduced target RNA cleavage was reported in the equivalent MjCsm2 Lys127Ala and AfCmr5 Lys148Ala mutants (65). However, mutation of the equivalent arginine residues in StCsm1 placed near sites 1 and 2 did not impact RNA cleavage (14). The required divalent metal ions could act to stabilize the developing negative charge at the transition state. Asp30, if it were present, could act as the general acid by donating a proton to the leaving oxygen. Asp30 may also participate in coordination of a divalent metal that enables a water molecule to donate the proton. Mutation of the strongly conserved Asp30-equivalent residue in several Csm3 homologs (Figure S8) was found to be deleterious to RNA cleavage. Since the putative general base arginine or tyrosine are supplied by Csm2 (sites 2-4) and Csm1 (site 1), Csm2 has a critical role, in addition to Csm3, towards achieving target RNA cleavage. This mechanism is consistent with the fact that 2’-deoxy substitution of the attacking oxygen (DTR) completely blocked target RNA cleavage (Figure 1E). DTR enhances the enzymatic activities of Csm1 (Figures 1F & 1G) through strengthening the interactions among the target RNA, the R and the H ridges or through multiple rounds of binding.

**Figure 5.**
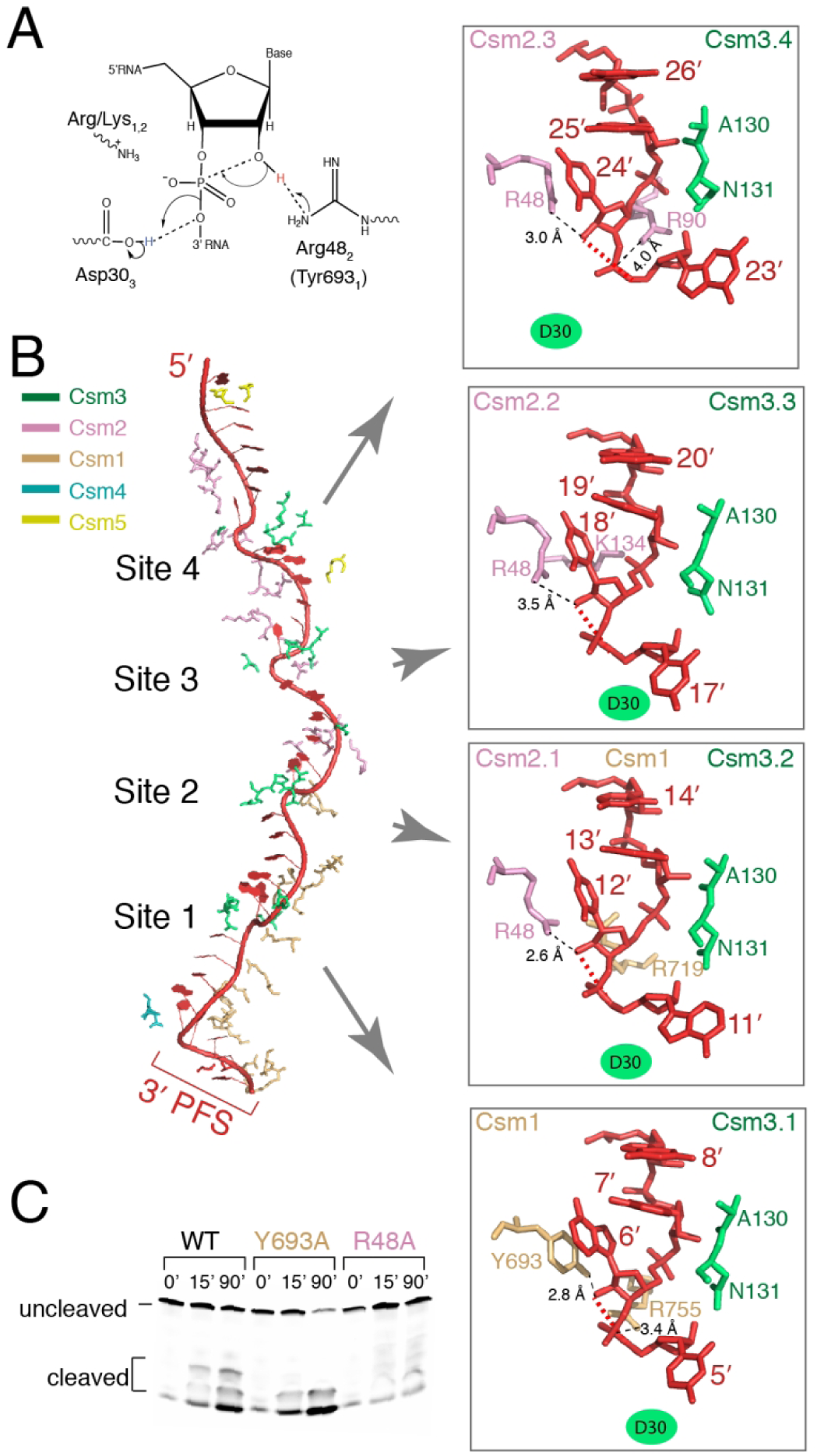
The structures of the four target RNA cleavage centers. (A). The RNase-A-like cleavage mechanism. (B) Structures of the four target RNA cleavage center. Each cleavage center is superimposed and displayed in the same orientation. Upon binding to the crRNA, target RNA undergoes base flipping every 6^th^ nucleotide at the four cleavage sites (Sites 1-4), forming an inline geometry involving the 2’-nucleophile oxygen, the scissile phosphate, and the 5’-leaving oxygen atoms. The residues of Csm1, Csm2 and Csm3 surrounding the cleavage sites are shown in stick models and labeled. The RNase catalytic site contained the Asp30Ala mutation in all solved structures. A possible reaction mechanism mediated by the conserved Csm3 (Asp30) and Csm2 (Arg48) residues is shown on top left. (C). In vitro RNA cleavage assay of the wild-type (WT), the Tyr693Ala (Y693A) of Csm1, and the Arg48Ala (R48A) mutants. The reaction products of 5’-Cy3-CTR for the wild type and mutants were analyzed at time points: 0 min, 15 min, 30 min, 60 min and 90 min.

### CTR Unleashes LlCsm1 from LlCsm4

The 3’ PFS is the key element of the CTR that activates Csm1 and is observed to form strategic interactions with Csm1. Four nucleotides, C(−1)’-A(−2)’-A(−3)’-G(−4)’, of the 3’ PFS in CTR make a 90 degree turn from the trajectory of the rest of the target RNA and rests on Csm1. Notably, three helices: α12 of the Palm 1 domain, α17 and α21 of the Palm 2 domain of Csm1 (Figure 2A & Figure S7) point their N-terminal ends towards the phosphate backbone of the 3’ PFS nucleotides, thereby forming a positive groove that interacts favorably with the negatively charged 3’ PFS phosphate backbone. The PFS nucleotides we used do not form any base-specific interactions with Csm1. Other 3’ PFS with different sequences could interact with Csm1 in a similar manner or to form additional interactions. The 3’ PFS-helix dipole interaction locks the two Palm domains in preparation for Csm1-mediated activities. Consistently, the target RNA lacking 3’ PFS (PTR) failed to induce either cOA synthesis or ssDNase activity of LlCsm1 (Figure 1F & 1G).

To unveil structural rearrangements, if any, upon CTR binding to LlCsm, we compared the apo structure (no target RNA) with the CTR-43 structure (cognate target RNA-bound). Within each complex, we computed and compared the transformation matrix of each subunit to its neighbor (Csm4 to Csm3.1, Csm3.1 to Csm3.2 and so on). Our results immediately revealed that subunits within the R ridge are related by an 8-fold screw symmetry to each other (∼45° rotation and 23 Å translation) in both the apo and the CTR-43 complexes (Figure S6). Interestingly in CTR-43, subunits within its H ridge also have the same 8-fold screw symmetry as well, suggesting that Csm3 and Csm2 have an inherent propensity to assemble into the same helical symmetry regardless if target RNA is bound (Figure S6). Furthermore, the 8-fold screw symmetry within RAMP protein assemblies was previously observed in both the crystal structure of Cmr4 and the Type III-B PfCmr cryoEM structure (20), suggesting a conserved structural property across Type III CRISPR-Cas subtypes.

The R ridges of both the apo and the CTR-43 complexes were superimposed to allow other subunits to be compared (Figure 3B). Not surprisingly, the bound crRNA, especially the portion that interacts with Csm3, superimposed well. Csm1 rotates ∼6 degrees upwards around an axis anchored just behind Csm4 from that in the CTR-43 to the apo model. Csm4 rotates the same amount, also upwards, but around a slightly differently anchored axis. As a result, Csm1 peals away from Csm4 and reduces its buried surface with Csm4 from an extensive 1403 Å^2^ in the apo to 1261 Å^2^ in the CTR-43 complex (Figures 2A, 3A & 3B).

In addition to rotation away from Csm4, Csm1 exhibits notable inter- and intra-domain movements upon CTR binding. While the Palm1 and the HD domains superimpose well, its Palm2 and D4 domains in CTR-43 are shifted upwards toward Csm2.1. As a result, the α21 helix (Csm1 residues 621-637) near the ATP-binding pocket moves upwards (Figure 3B), which could allow ATP to enter the GGDD active site and be converted to cOA.

Other structural elements in Csm1 and Csm4 previously discussed underwent minimal changes. To highlight these key structural elements, we assigned common names to two Csm1 loops (Figure S7) and one Csm4 loop (Figure S9). The Linker-Palm1 (LiP) loop (LlCsm1 residues 394-416) and the LASSO loop (LlCsm1 residues 86-103) superimpose very well between the apo and the CTR-43 structures (Figure 3B). Note that the well conserved zinc-ribbon motif of Csm1 resides at the end of LiP (C402, C405, C418 and C421) (Figure S7) that was predicted to play an important role in target RNA binding (66). Interestingly, mutation of the corresponding StCsm LiP loop (StCsm1 residues 393-417) severely impaired and that of the previously studied *Thermococcus onnurineus* Csm, ToCsm, LASSO loop (ToCsm1 residues 107-115) constitutively activated Csm1 DNase activities (13, 14), suggesting conformational changes in these loops are not the causes for activation. Similarly, the Csm4 Lid loop (Csm4 residues 82-96) also showed non-detectable structural changes between the two structures. Thus, the minimal structural change observed in the HD domain upon CTR binding are unlikely to explain the activation of the DNase activity.

### 3’ PFS of NTR Stabilizes LlCsm1 onto LlCsm4

Unlike CTR, NTR binding leads to inhibition of both the DNase and cOA synthesis activities in LlCsm (Figure 1F and Figure 1G). Surprisingly, regions in LlCsm1 that undergo changes from apo to CTR-43 also occur in NTR-bound LlCsm, leading to a high structural similarity between the NTR- and CTR-bound LlCsm1 (R.M.S.D 1.3 Å for 5984 atoms) (Figure 4). The α21 helix in Palm 2 domain in the NTR-bound LlCsm shows a similarly large upward motion from the apo (Figure 4B) as α21 of CTR-43 (Figure 3B). Similarly, the D4 domain of Csm1 in the NTR-bound complex also rests at a similar location as that in CTR-43 (Figure 4B). Thus, the remarkable difference in enzyme activity observed between the NTR- and the CTR-bound LlCsm cannot be explained simply by Csm1 structural change.

The 3’ PFS of NTR differs from that of CTR-43 and bears complementary sequence to that of the crRNA 5’ tag. Seven of the eight PFS nucleotides are observed and only four form base pairs with the 5’ tag (Figure 4). Nucleotide (−1)’ is flipped out by the thumb loop of Csm4 similar to those at the cleavage sites. Nucleotides (−2)’-(−5)’ base pair with nucleotides (2-5) of the 5’ tag and (−6)’ -(−7)’ rest on a groove formed by the Lid loop of Csm4 and the LiP loop of LlCsm1 (Figure 4). Though the Csm1-Csm4 interface further reduces from 1261 Å^2^ in the CTR-bound to 929 Å^2^ in the NTR-bound complex, LlCsm1 gains 433 Å^2^ interface with 3’ PFS that in turn bases pairs with the 5’ tag of crRNA (Figure 4B). As a result, 3’ PFS establishes an extensive interface that stabilizes Csm1 onto Csm4 and crRNA.

### Structural Dynamics of Csm1 and Csm2 Underlies LlCsm Activity

The differential interaction between Csm1 and Csm4 among the CTR-bound, the NTR-bound and the apo complex suggests that Csm1 has different stability but similar structure in different complexes. We thus examined the carefully refined atomic displacement parameter, or the B factor, combined with the map-fitting quality score, Q score (58), to quantify the dynamic behavior of each residue. Under the assumption of unit occupancy and isotropic motion, B factors are atomic displacement parameters of the Gaussian distribution of positions and thus can measure the degree of motion (67). Since B factors can be artificially high in areas with weak density, we also computed Q-scores, which reflect the map fitting quality independent of the B factors (58). As shown in Figure 6, while LlCsm1 of the apo complex has a relatively uniform distribution of B factors, that of the CTR-43 has a steep gradient with the HD domain reaching the highest B factors, reflecting a high mobility in this region (Figure 6A). Interestingly, accompanying the increase in motion of LlCsm1 is the low B factors or rigidification of the Csm2-based H ridge (Figure 6A), suggesting that the increase in Csm1 entropy is in part at the expense of the Csm2 rigidification, consistent with its structural role in binding and cleaving the target RNA. In both Csm2-bound complexes, the Csm3-based R ridge have similarly low B-factor values (Figure 6A), which reflects a similarity in dynamics of these regions. The computed high Q scores indicate good model fitting throughout each of the complexes except for a decrease in the HD domain of CTR-43 owing to the lower local resolution (Figure 6B).

**Figure 6.**
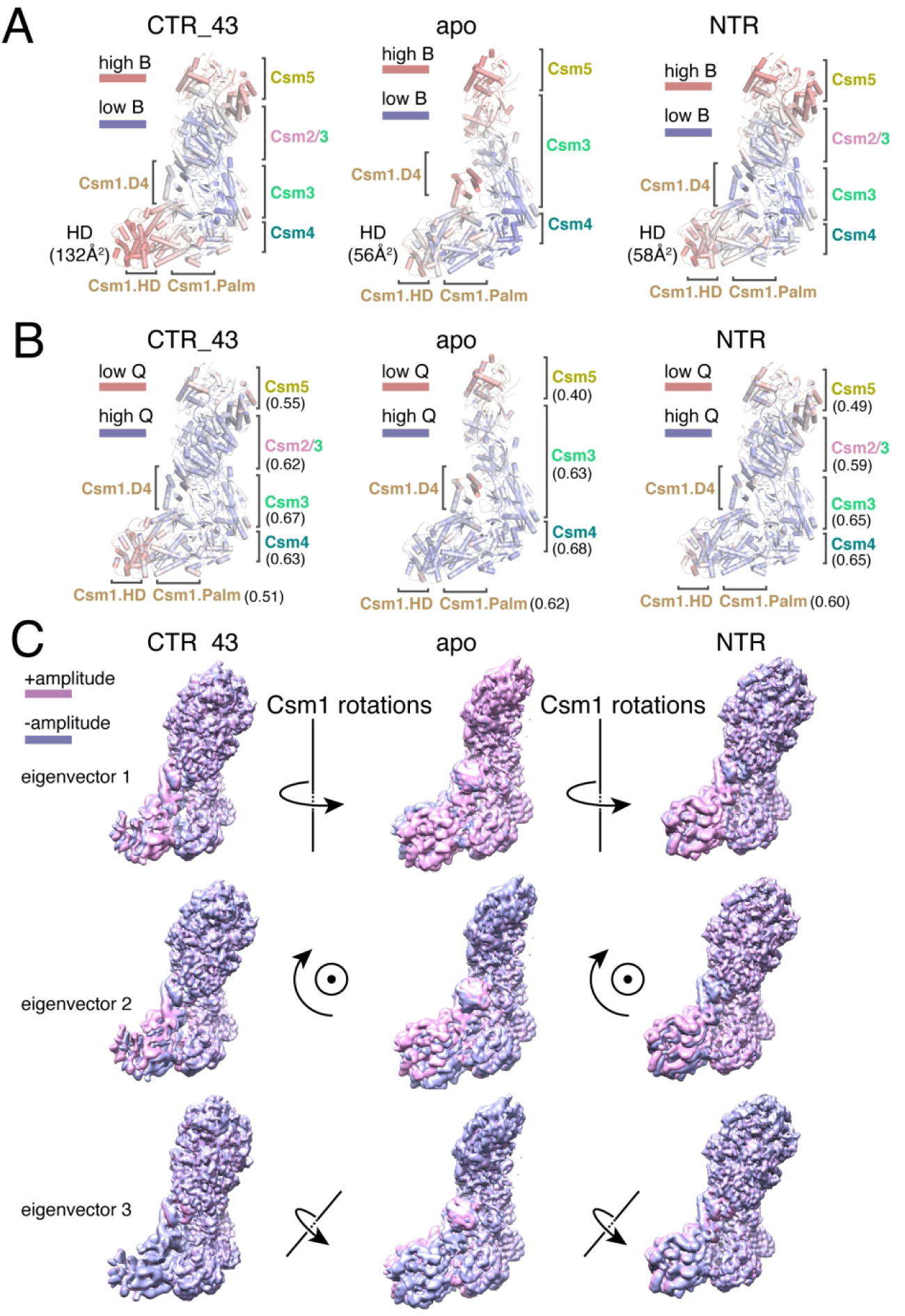
Dynamic features in the apo, the CTR-43 and the NTR complexes. (A) B factors of each complex are plotted on the refined models in a color gradient with blue being low and red being high values. Location of the subunits or domains of Csm1 are labeled and the average B factor of the HD domain is included in the parenthesis for each complex. (B) Q scores of each complex are plotted on the refined models in a color gradient with blue being high and red being low values. Location of the subunits or domains of Csm1 are labeled. Average Q-scores for all subunits are included in the paranthesis. (C) 3D multivaribilty analysis for the LlCsm complexes. Electron potential densities representing the two extreme amplitude values of the top 3 eigenvectors are superimposed at the Csm3 subunits and displayed as pink and blue, respectively. The direction of the motion for each eigenvector is indicated.

To better understand the conformational landscape of the mobile Csm1 subunit, we performed multi-body refinement on all three complexes with RELION-3 (48). The multi-body analysis uses the three pre-defined rigid body shapes to simultaneously align each rigid body projection while subtracting the other two. The process is repeated for all three rigid bodies, thus resulting in relative orientation for each body in each particle. We defined three rigid bodies: Csm5, Csm4, Csm3, and Csm2 defines the first, the HD domain of Csm1 defines the second and the rest of Csm1 defines the third. We analyzed the continuous structural heterogeneity with regards to these three bodies present in the samples. To our surprise, although Csm1 is clearly moving in all three samples, it has very similar motions, which does not completely explain the difference in the B factors among the three complexes. We interpret this result as that Csm1 has similar intrinsic motions in all states, as indicated by the similar direction and magnitude of the orthogonal set of motions, or eigenvectors (Figure 6C & Supplementary Movie S1-S3). However, the type of dynamic motion induced by CTR may be different, at a fast time scale or not describable by rigid-body motions, for instance, from its inherent motion measured by the 3D variability analysis. The fact that the CTR-bound complexes have the high mobility in Csm1 and low mobility in Csm2 and vice versa in the two inactive complexes suggests a correlated dynamic change throughout the LlCsm complex with the enzymatic function.

### Csm2 Deletion is Detrimental to LlCsm Activities

The dynamic model predicts an important role of Csm2 in LlCsm function. To assess how Csm2 impacts the enzymatic activities associated with Csm1, we purified a mutant LlCsm whose Csm2 subunit is deleted (ΔCsm2-LlCsm) (Figure 7A) and compared its enzymatic activities with those of the intact LlCsm complex. As expected, ΔCsm2-LlCsm complex has significantly reduced target RNA cleavage activity (Figure 7B). Removal of Csm2 also weakened CTR-activated phage DNA cleavage (Figure 7C) and cOA synthesis activities while DTR-activated overnight cOA synthesis remained intact, likely due to multiple rounds of activations (Figure 7D).

**Figure 7.**
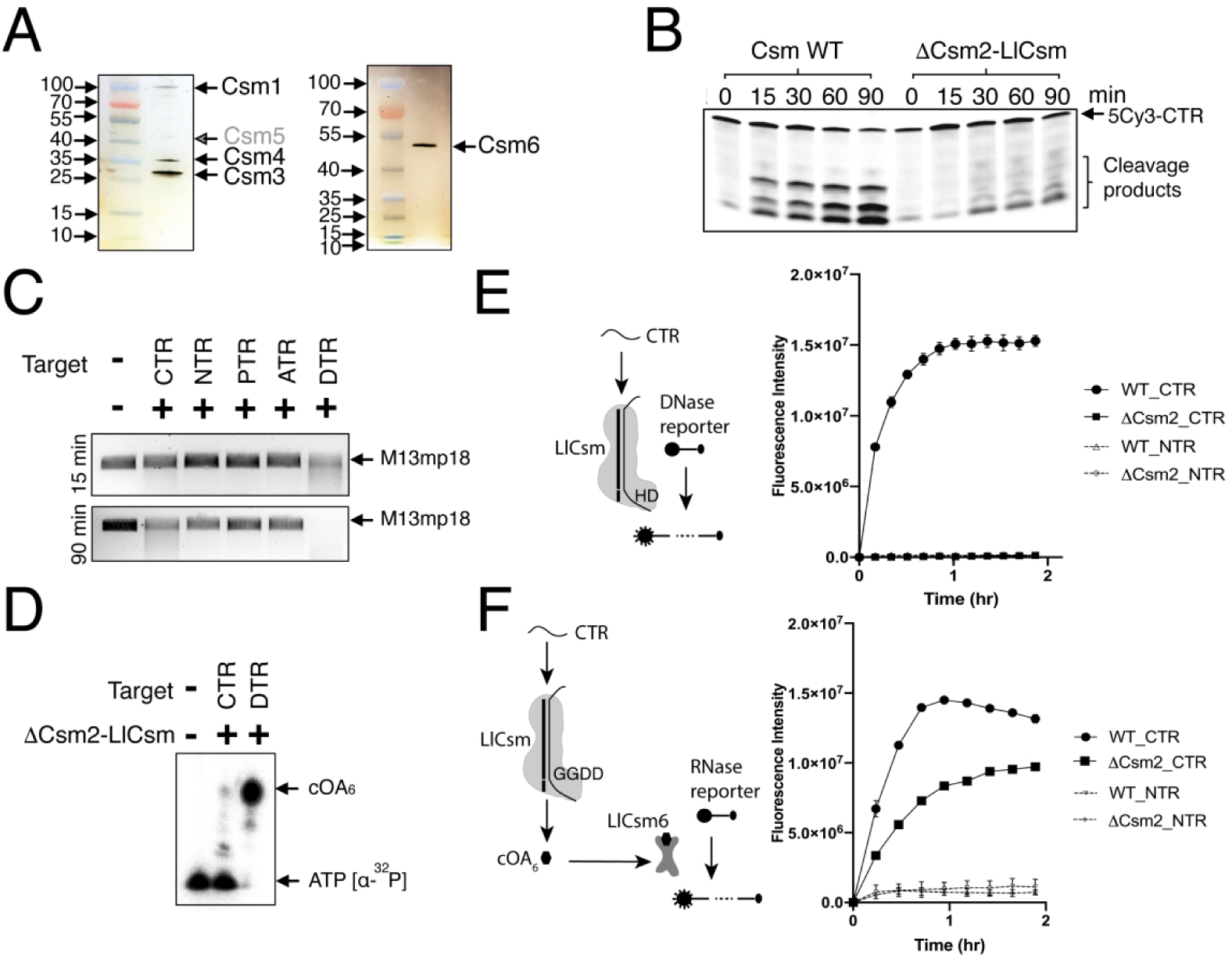
Enzymatic activity comparison between the wild-type (WT) and LlCsm without Csm2 (ΔCsm2-LlCsm). (A) Silver stain profiles of the co-purified ΔCsm2-LlCsm complex (left) and His-tagged LlCsm6 protein (right). The ΔCsm2-LlCsm complex comprises of intact Csm1, Csm4 and Csm3. Csm5 that copurified with His-tagged Csm3 is low but present. (B) *In vitro* RNA cleavage assay by the WT and ΔCsm2-LlCsm complex. The RNase reaction products of 5’-Cy3-CTR were analyzed at time points: 0 min, 15 min, 30 min, 60 min and 90 min. (C) *In vitro* DNA cleavage assay by the ΔCsm2-LlCsm complex. M13mp18 was used as substrate and the reaction products were analyzed at time points 15 min and 90 min. (D) *In vitro* cOA synthesis assay by the ΔCsm2-LlCsm complex. Radiolabeled α-^32^P-ATP was used as a substrate and the reactions were incubated overnight. (E) Schematic and the results of a fluorescence DNase reporter assay that reports CTR-induced and Csm1-HD domain-mediated DNA cleavage. The rise in fluorescence intensity (arbitrary units) indicates cleavage by Csm complex. Triplicates were performed for each indicated reaction. (F) Schematic and results of a fluorescence RNA reporter assay that reports CTR-induced, Csm1-GGDD motif-mediated cOA synthesis and cOA-stimulated LlCsm6 cleavage of RNA. The rise in fluorescence intensity (arbitrary units) indicates production of cOA by Csm that stimulated RNA cleavage by Csm6. Triplicates were performed for each indicated reaction.

To better dissect the kinetic process of DNase and cOA synthesis of LlCsm in the presence and absence of Csm2, we performed two fluorescence-based activity assays. The DNase activity is detected by the rise in fluorescence intensity of a labeled DNA probe upon its cleavage by LlCsm (Figure 7E). The wild type LlCsm showed a significant increase in fluorescence over time upon addition of CTR but not NTR (Figure 7E). In case of ΔCsm2-LlCsm, however, CTR elicits minimal fluorescence over time (Figure 7E), consistent with a requirement of Csm2 for the CTR-activated DNase activity of LlCsm.

The cOA synthesis activity is indirectly measured by the rise in fluorescence intensity of a labelled RNA probe upon its cleavage by LlCsm6 whose ribonuclease activity is stimulated by LlCsm-produced cOAs in the same reaction (Figure 7F). The cleavage of RNA probe, and consequently the rise in fluorescence, therefore, reflect the activity of LlCsm6 coupled to cOA synthesis by LlCsm. Consistently, addition of CTR but not NTR to a reaction mixture containing LlCsm, LlCsm6, and ATP induced high fluorescence (Figure 7F). Interestingly, CTR also induced notable fluorescence increase when added to the reaction mixture containing ΔCsm2-LlCsm, LlCsm6 and ATP, although at a slower rate (Figure 7F), suggesting that ΔCsm2-LlCsm still supports the synthesis of cOA, although with a notable delay, which resulted slower RNA cleavage by LlCsm6. Together with weakened DNase activity in ΔCsm2-LlCsm, these results support a direct impact of Csm2 on Csm1 catalytic functions.

Finally, we tested ΔCsm2-LlCsm activity in the plasmid interference. Strikingly, ΔCsm2-LlCsm was found to have a profound negative effect on the plasmid interference activity (Figure S2, Table S1). Similar to the Palm2 deficient LlCsm, ΔCsm2-LlCsm was not able to interfere with the target plasmid, suggesting an important role of Csm2 in regulation of Csm1-mediated activities in cells.

## Discussion

The Type III-A, or Csm, complex is a multifaceted immune effector owing to its three separate enzymatic activities and possesses a remarkable mechanism of enzymatic activity control. Initially triggered by transcribing viral RNA, Csm unleashes its RNase, ssDNase, and cOA_n_ synthesis activities. Subsequently, cleaved RNA dampens the latter two activities in order to limit the immune response to a narrow temporal window. Self RNA, which could occur if host CRISPR arrays were transcribed in the antisense direction, differs from viral RNA at its 3’ flanking sequence and is able to base pair with the 5’-tag sequence of the crRNA. Interaction of Csm to self RNA (NTR) switches off both ssDNA and cOA_n_ synthesis activities and protects the host cell from potentially lethal autoimmunity. Although not related to Csm regulation, cOA_n_ goes on to elicit secondary immune responses. Our current understanding of the molecular basis responsible for the remarkable Csm regulation mechanism remains incomplete.

The enzymatic activities of Csm1 depend on its assembly status. In either isolated (ssDNase) or the CTR-bound state, Csm1 is active while in the apo or the NTR-bound state, Csm1 is inactive. Exhaustive structural comparison of StCsm(14), ToCsm(13), and now LlCsm (this study), at different states reveal no common molecular transitions among the four states that can explain Csm1 activation. The large sliding of the Csm2-formed H ridge upon target RNA binding is shared between both the inactive NTR-bound and the active CTR-bound states. Local rearrangements within LiP or LASSO from the NTR- and the CTR-bound transition have different structural characteristics among the three Csm complexes (13, 14). Notably, despite being distant from the active and unconserved, mutations of LiP, L1 or LASSO do impact Csm1 activities (13, 14), suggesting that they contribute to changes of a physicochemical property of Csm1 critical to its enzymatic activities. These studies, then, proposed that dynamics underlies activation. The first experimental evidence supporting a dynamics-based activation model was presented in a single-molecule fluorescence microscopy study (26). It showed that CTR triggers fast conformational fluctuations within Csm1 while NTR locks it in a rigid conformation (26). However, the molecular basis of how the two different target RNA change the dynamics of Csm1 was not elucidated by single molecule studies.

Our results that the histag-Csm2-based H ridge is missing from the inactive apo LlCsm structure clearly indicates its weak association with the complex or of large dynamic motions. The correlated change in Csm1 dynamics with that of Csm2 and its enzymatic activity is striking. Accompanied by the weak binding of Csm2, and to some extent, of Csm5, Csm1 is well ordered in the apo complex and locked in an inactive state. In contrast, Csm2 is well ordered in the CTR-bound complex while Csm1 is in a relaxed state (Figure 6), which enables its ssDNA cleavage and cOA_n_ synthesis activities. In the presence of NTR, Csm2 is present but has elevated motions than in the CTR-bound state and Csm1 remains rigid. In this case, 3’PFS of the NTR forms a completely different interaction than that of CTR that enhances Csm1 rigidity through its interaction with Csm4. While the dynamic state of Csm1 observed by cryoEM is consistent with that by the single-molecule fluorescence study (26), we further observed a dynamic correlation between Csm2 and Csm1, and among the other structural elements in LlCsm that best explains the regulation of its activity than conformational changes.

Deletion of the Csm2 subunit from the effector complex was found to be detrimental to the *in vivo* plasmid interference activity, target RNA cleavage, and ssDNA cleavage, although maintained some *in vitro* cOA synthesis capabilities. The importance of Csm2 to *in vivo* immunity reflects its observed structural role in balancing the dynamics of LlCsm in its activity. Previously, Csm6-mediated RNase activity was implicated to be more relevant to achieve anti-plasmid immunity than Csm1-mediated DNase activity (60). Consistently, we observed complete shutdown of plasmid interference upon mutating the Csm1 GGDD motif to GGAA (Figure S2), suggesting that cOA synthesis and consequently, Csm6 activation, is required to achieve anti-plasmid immunity in this system and that Csm2 can impact cOA synthesis. Recently, a novel type III-A *Lactobacillus delbrueckii* subsp. *bulgaricus* (LdCsm) complex that lacks any detectable Csm6 homologs and harbors a cOA-synthesis deficient QGDD motif unlike the consensus GGDD motif (Figure S7) was found to mediate anti-plasmid interference in *E.coli* (68). This suggests a possibility that the Csm DNase activity plays a secondary role in immunity but can still defend invading nucleic acids in absence of cOA-mediated Csm6 activity. In this case, Csm2 could play a role in maintaining the Csm DNase activity.

Previous studies of *Streptococcus thermophilus* Csm and *Thermococcus onnurineus* Csm identified regions of Csm1 that are linked to its activation (13, 14). As these regions are scattered through Csm1 and are not physically close to the active sites, we suggest that these data can be united under the protein dynamic regulation model. Inhibition of the StCsm1 LiP loop mutants to StCsm activities can be explained in part by their inability to induce the necessary dynamic motion required for the activity. In the case of the LASSO loop, that when mutated, led to constitutive activation of Csm1, the Csm complex may undergo a separation between the HD domain and the Palm1/2 domains, leading to the increased dynamics, thus activity.

The protein dynamics-mediated activation model also explains why Csm1 alone (31) is active in ssDNase cleavage. In this case, the HD domain is not restrained and thus active. In contrast, when Csm1 is bound within the apo Csm RNP, it is immobilized in an inactivate state. Further testing of this activation model awaits additional computational and other biophysical studies.

## Author contributions

The LlCsm plasmid was provided by M.P.T to H.L. S.S performed all aspects of cryo-EM sample preparation: purification of effector complexes and complexing with target RNA. J.R and S.S. prepared cryoEM grids. J.R. performed data collection. J.R. and H.L. performed data processing, model building and structure refinement. S.S. and C.W. performed expression and purification of variants of LlCsm effector and LlCsm6 protein. W.W. performed *in vivo* plasmid interference assays. S.S performed *in vitro* RNase, ssDNase and fluorescence reporter assays. C.W. performed cOA synthesis assays. H.L. and S.S. wrote the manuscript. S.S., J.R., W.W., M.P.T. and H.L. made the figures. All authors edited the manuscript. No conflict of interest declared.

## Acknowledgments

The authors wish to acknowledge and thank the valuable support of the following personnel working at Florida State University: Xiaofeng Fu, Biological Science Imaging Resource (BSIR), Brian Washburn and Cheryl Pye, Molecular Cloning Facility, and Steve Miller, DNA Sequencing Facility. The authors acknowledge the use of instruments at the Biological Science Imaging Resource of Florida State University. Its Titan Krios was purchased in part from NIH grant S10 RR025080, the Vitrobot and Solaris Plasma Cleaner from S10 RR024564 and the BioQuantum/K3 from U24 GM116788. This work was supported by NIH Grant R01 GM099604 to H.L. and NIH Grant R35GM118160 to M.P.T.

## Data Deposition

CryoEM maps (coordinates) have been deposited to Protein Data Bank with identification codes EMD-22266 (6XN3), EMD-22267 (6XN4),EMD-22269 (6XN7), EMD −22268 (6XN5) for the CTR-43 complex, the CTR-32 complex, the NTR complex, and the apo complex, respectively.

## SUPPLEMENTARY INFORMATION

**Figure S1.**
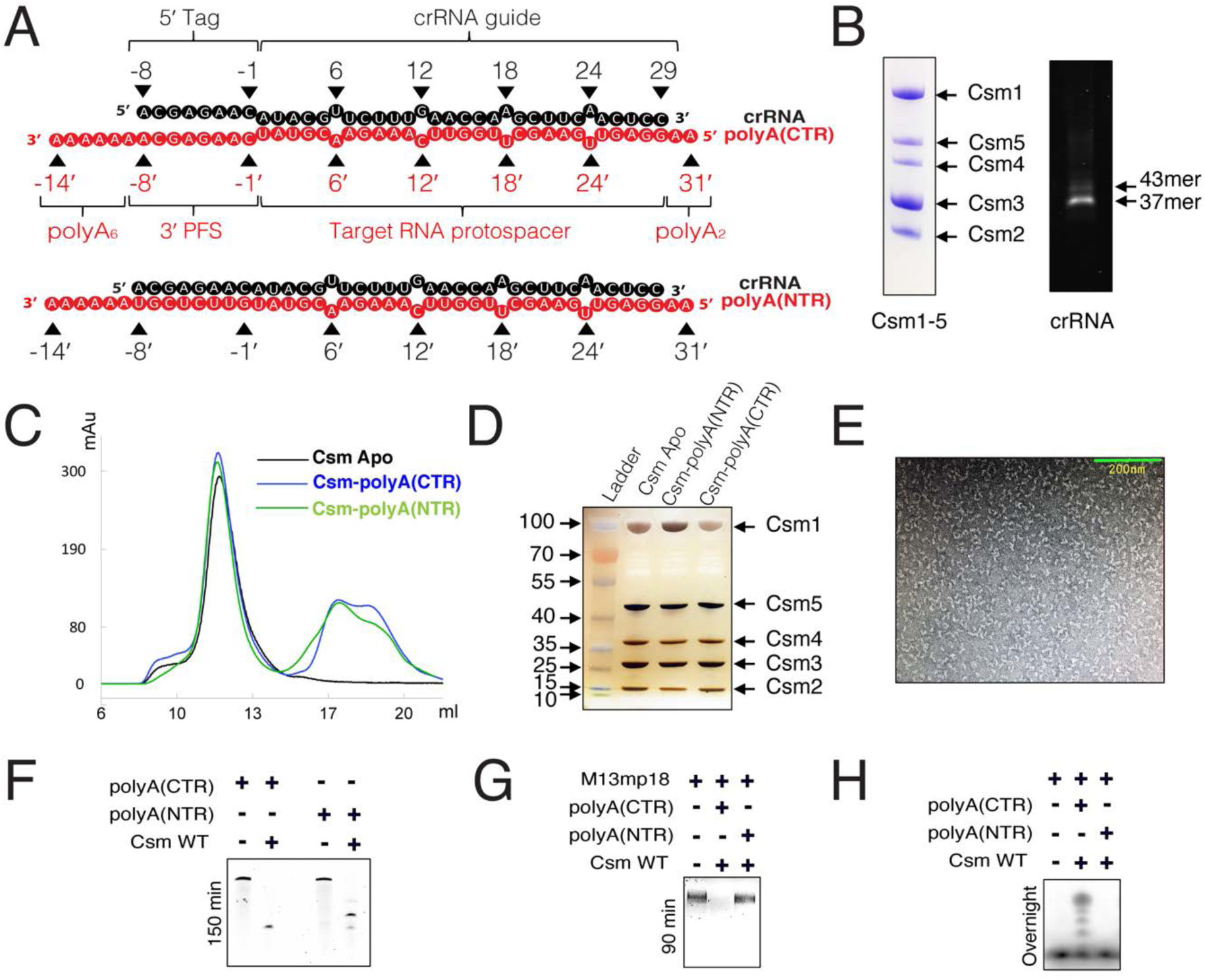
Cryo-EM sample preparation and activity tests on target RNA. Related to Figures 1, 2, 3 and 4. (A) Schematic representation and nomenclature of crRNA-polyA(CTR) and crRNA-polyA(NTR) duplexes. The target RNAs used in the cryo-EM samples contained 6-nt and 2-nt polyA extensions at 3’- and 5’-ends respectively. (B) Sodium dodecyl sulfate - polyacrylamide gel electrophoresis (SDS-PAGE) and denaturing Urea PAGE gel images showing the quality of *L. lactis* ribonucleoprotein (RNP) used to make target-free and target-bound complexes for structural studies. The purified RNP contained all five proteins forming the effector complex (Csm1-5) and two species of crRNA (major 37mer species and minor 43mer species). (C) Overlay of size-exclusion chromatograms of no-target (apo), polyA(CTR) and polyA(NTR)-bound Csm complexes in black, blue and green colors respectively. (D) Silver stain profiles of no-target (apo), polyA(CTR) and polyA(NTR)-bound Csm complexes showing the presence of all 5 Csm subunits in all sample preparations. The loaded amounts are too low for crRNA to be visible. (E) Negative stain showing homogenous distribution of LlCsm particles during sample screening. The image was collected using FEI CM120 Biotwin transmission electron microscope. (F) *In vitro* RNase cleavage assay showing that polyA(CTR) and polyA(NTR) used for structure determination is similar to RNase activity by Csm WT on CTR and NTR respectively. (G) *In vitro* DNase cleavage assay showing that polyA(CTR) and polyA(NTR) used for structure determination is similar to DNase activity by Csm WT on CTR and NTR respectively. (H) *In vitro* cOA synthesis assay showing that polyA(CTR) and polyA(NTR) used for structure determination is similar to cOA synthesis by Csm WT on CTR and NTR respectively.

**Figure S2.**
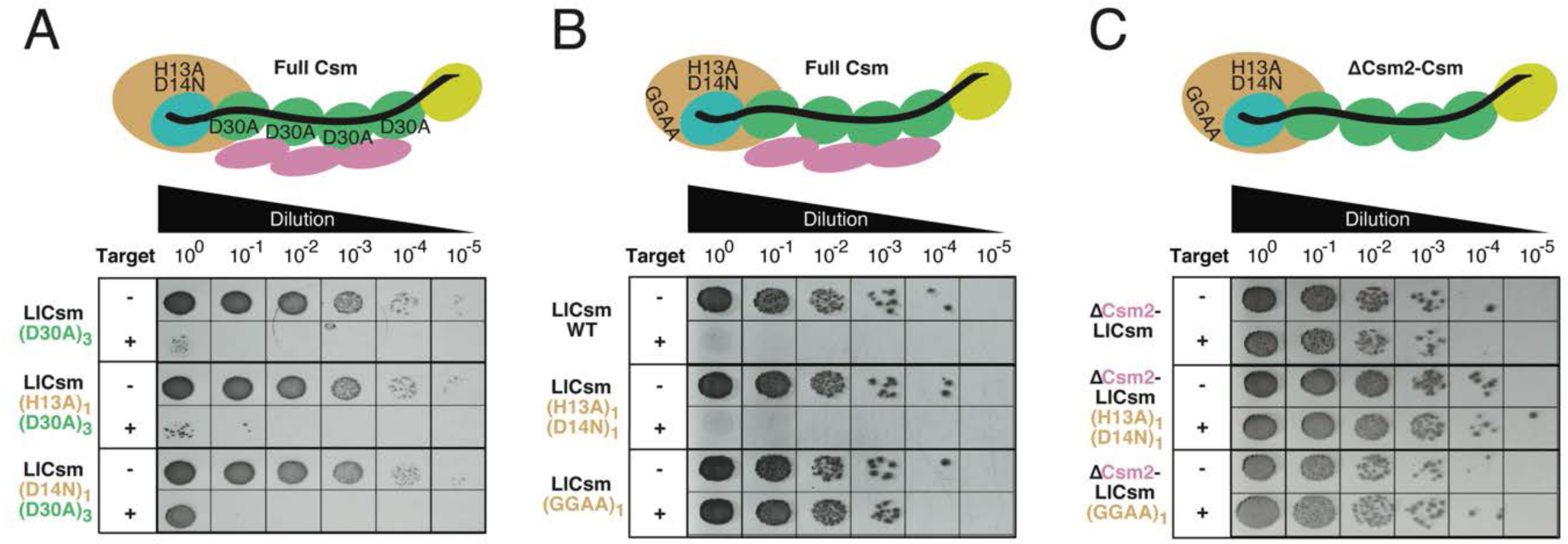
Assessment of *in vivo* functionality of LlCsm complex using plasmid interference assay in bacterial cells. Related to Figures 2, 3 and 7, Table S1. (A) Plasmid interference activities of full LlCsm complex harboring Csm1 and/or Csm3 mutations. (B) Plasmid interference activities of full LlCsm harboring Csm1 HD mutation or Csm1 palm2 mutation. (C) Plasmid interference activities of ΔCsm2-LlCsm harboring Csm1 HD mutation or Csm1 palm2 mutation.

**Figure S3.**
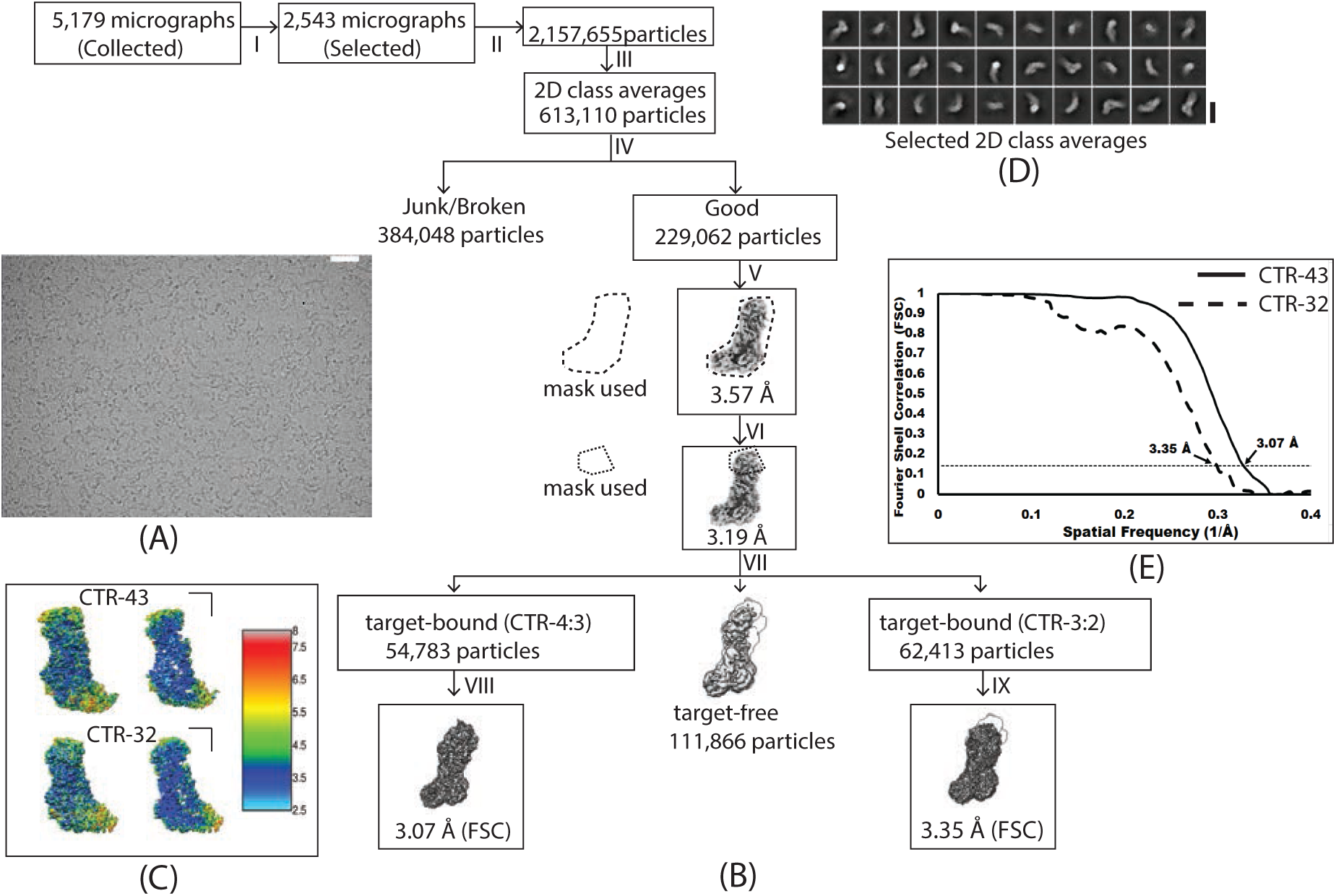
Cryo-EM data processing and refinement flow chart of LlCsm-CTR-43 and LlCsm-32 complex. Related to Figure 2. (A) Raw micrograph (scale bar 20 nm) (B) Image processing flowchart: In total 5,179 micrographs were collected. 2,543 were chosen for the frame alignment. 2,15,655 particles were extracted using Relion 3.0 (1) and imported into cryoSPARC software (2) to perform multiple rounds of 2D analysis. After 2D analysis, 613,110 particles were selected and imported to Relion 3.0 to perform 3D classification that resulted in 229,062 particles which further refine to 3.57 Å resolution and further per particles CTF refine followed by autorefine improved the resolution to 3.19 Å. 3D classification was performed using mask on the top part (Csm5), which sorted out the particles into three classes. First class contain four copies of Csm3 densities and three copies of Csm2 densities (CTR-43). Second class lack Csm2 densities and target RNA densities. Third class contain the three copies of Csm3 densities and two copies of Csm2 densities (CTR-32). 54,783 particles from first class was imported into CisTEM software (3) and further refined to obtain 3.07 Å resolution. 62,413 particles from third class were further refined using per particle CTF refine and autorefine that resulted in 3.5 Å. (C) Local resolution estimated by Resmap. (D) Selected 2D class averages (scale bar 15 nm). (E) FSC curves.

**Figure S4.**
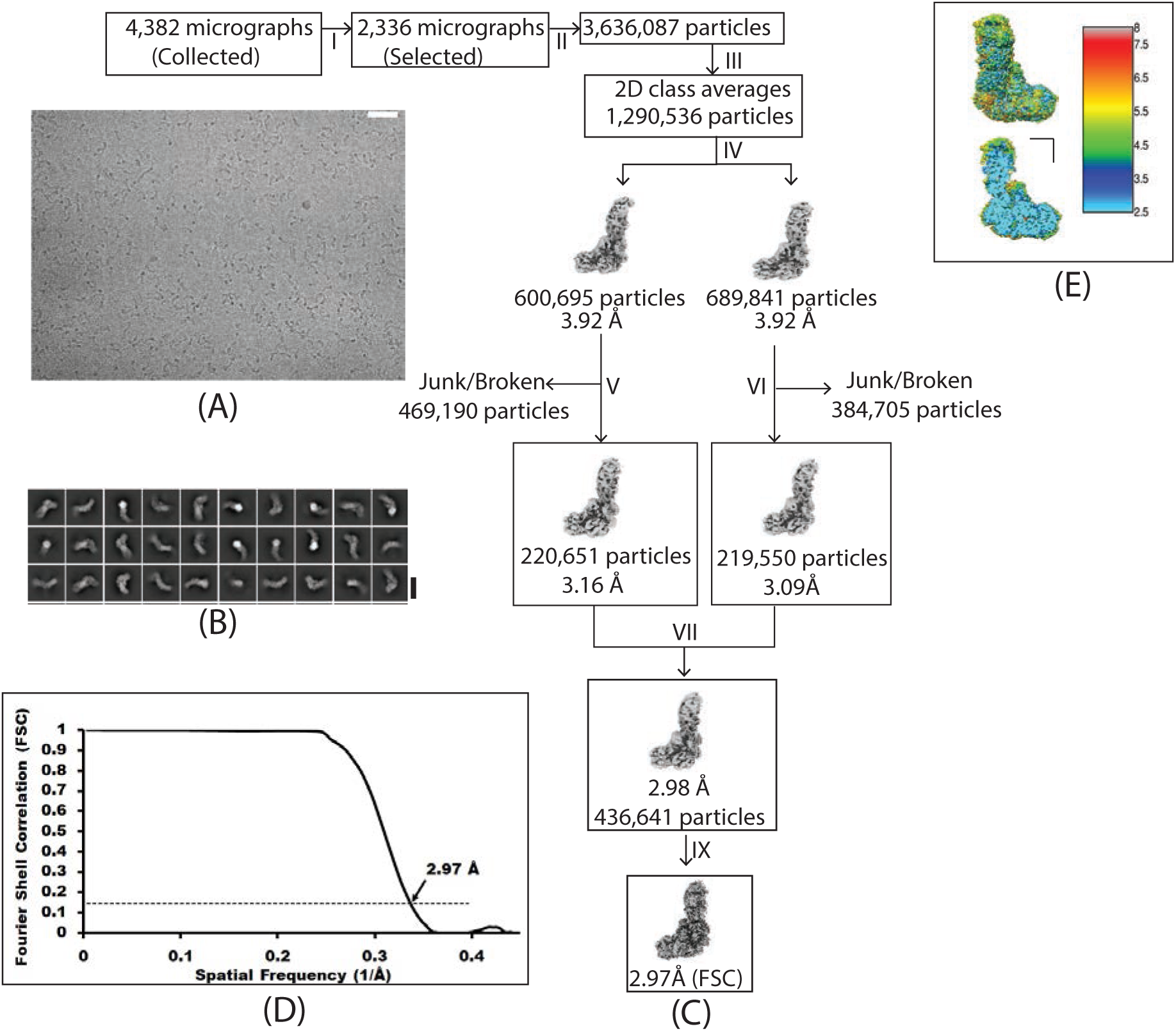
Cryo-EM data processing and refinement flow chart of LlCsm-apo complex. Related to Figure 3. (A) Raw micrograph (Scale bar 50 nm). (B) Local resolution estimated by Resmap. (C) Image processing flow chart: In total 4,382 micrographs were collected. 2,336 were chosen for the frame alignment. 3,636,687 particles were extracted using RELION 3.0 (4) and imported to cryoSPARC software (2) to perform multiple rounds of 2D analysis. After 2D analysis, 1,290,536 particles were selected and imported to RELION 3 (1) to perform 3D classification that resulted in 436,641 good particles which were used for per particle CTF refine followed by autorefine yielding the resolution of 2.98 Å. Furthermore, the particles were imported in cisTEM which resulted in the resolution of 2.97 Å. (D) Selected 2D class averages (scale bar 15 nm). (E) FSC curve.

**Figure S5.**
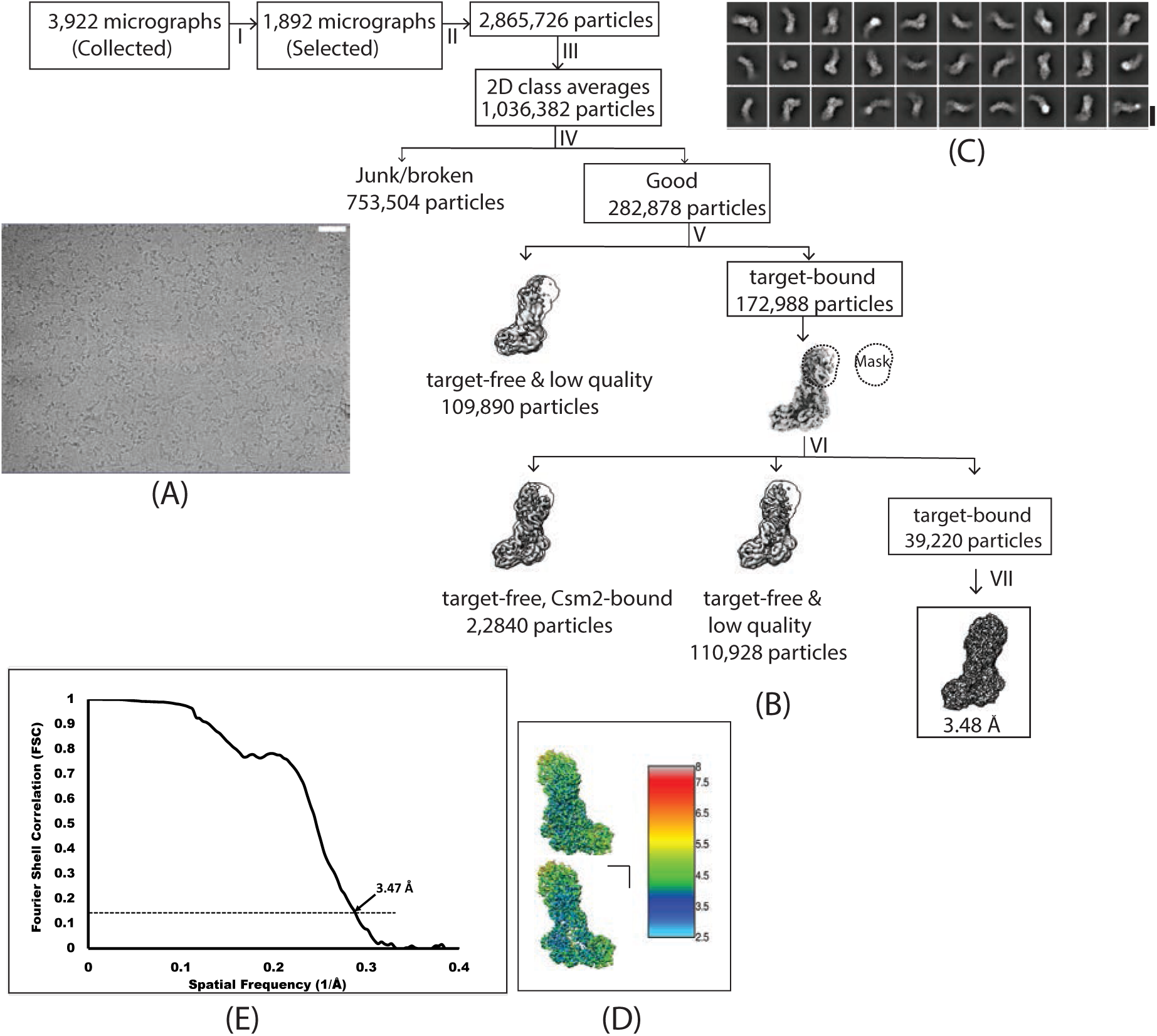
Cryo-EM data processing and refinement flow chart of LlCsm-NTR complex. Related to Figure 4. (A) Raw micrograph (Scale bar 50 nm) (B) Image processing flow chart: In total 3,922 micrographs were collected. 1,892 were chosen for the frame alignment. 2,86,726 particles were extracted using RELION 3.0 (4) and imported to cryoSPARC software (2) to perform multiple rounds of 2D analysis. After 2D analysis, 1,036,382 particles were selected to perform 3D classification that resulted in 282,878 particles followed by another round of 3D classifications. The particles with target RNA were pooled together and imported into RELION for classification based on Csm5 using mask on the top part (Csm5) resulting in three classes. The two classes showed no density of target RNA and were discarded. The third class with target density was further refined with ctf refine that resulted in 3.48 Å resolution. (C) Local resolution estimated by Resmap. (D) Selected 2D class averages (scale bar 15 nm). (E) FSC curve.

**Figure S6.**
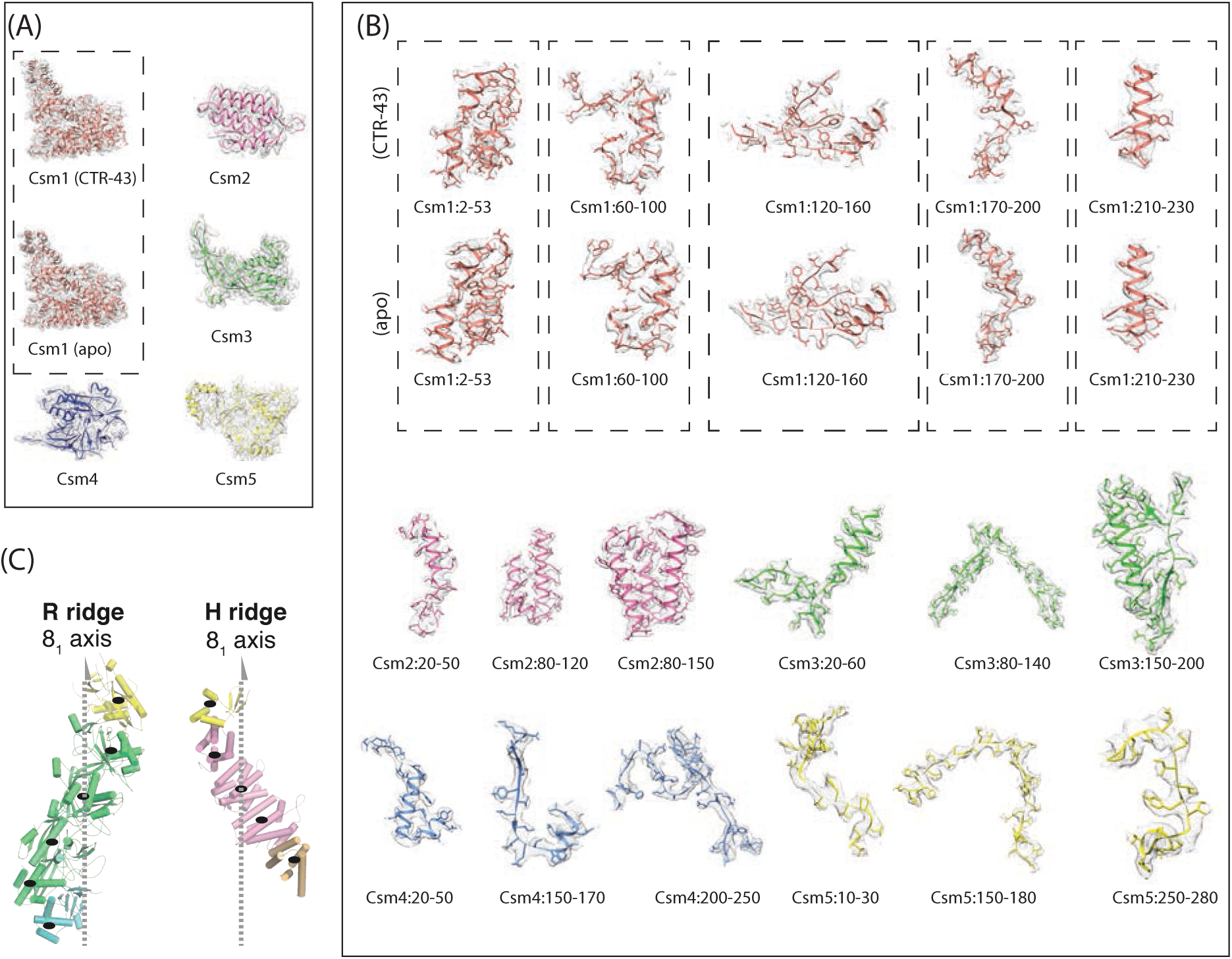
Density maps superimposed with finally refined models. Related to Figures 2, 3 and 4. (A) The density map of each subunit is from the CTR-43 structure while that of Csm1 from the apo structure is also shown for comparison. (B) Focused views of the density for selected regions are shown. The densities of Csm1 are shown for both CTR-43 and apo structures for comparison. (C) Isolated R and H ridges with the 8-fold screw axis indicated.

**Figure S7.**
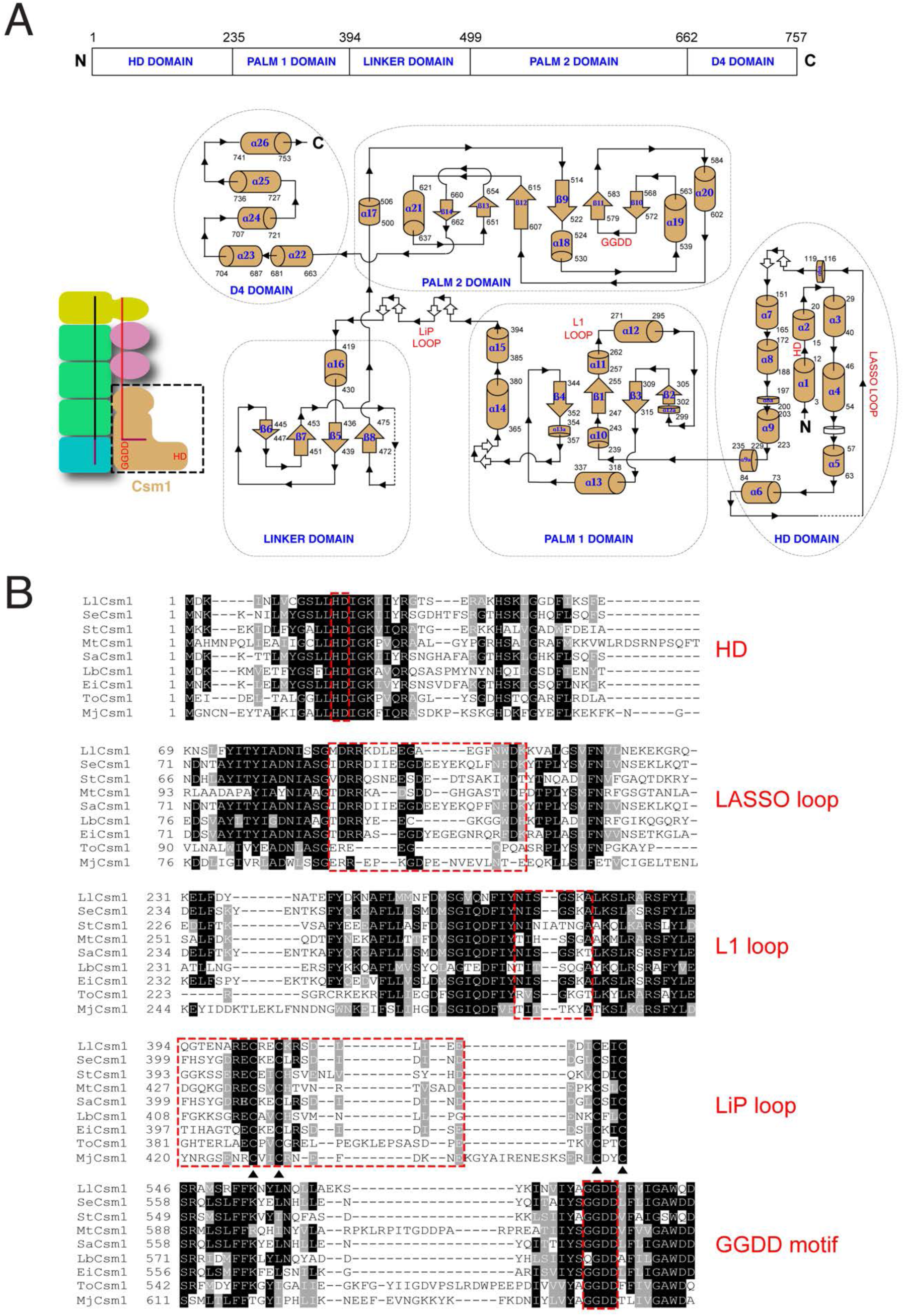
Domain organization, topology and multiple sequence alignment of LlCsm1. Related to Figures 2, 3 and 4. (A) Topology map of LlCsm1 composed of N-terminal HD domain, palm1 domain, linker domain, palm2 domain and C-terminal D4 domain. The DNase HD catalytic site, the GGDD motif constituting the cOA synthesis site and the loops important in dynamic functioning: Lasso loop, L1 loop and LiP loop are highlighted. (B) Multiple sequence alignment of *Lactococcus lactis* Csm1 (LlCsm1) with bacterial Csm1 orthologs from *Staphylococcus epidermidis* (SeCsm1), *Streptococcus thermophilus* (StCsm1), *Mycobacterium tuberculosis* (MtCsm1), *Staphylococcus aureus* (SaCsm1), *Lactobacillus delbrueckii* subsp. *bulgaricus* (LbCsm1), *Enterococcus italicus* (EiCsm1) and archaeal orthologs from *Thermococcus onnurineus* (ToCsm1), *Methanocaldococcus jannaschii* (MjCsm1). The red dashed boxes highlight highly conserved HD residues, sequence-variable lasso loop, L1 loop, LiP loop and conserved GGDD motifs. The black triangles indicate four strongly conserved cysteine residues forming the Zinc-finger motif in Csm1 orthologs.

**Figure S8.**
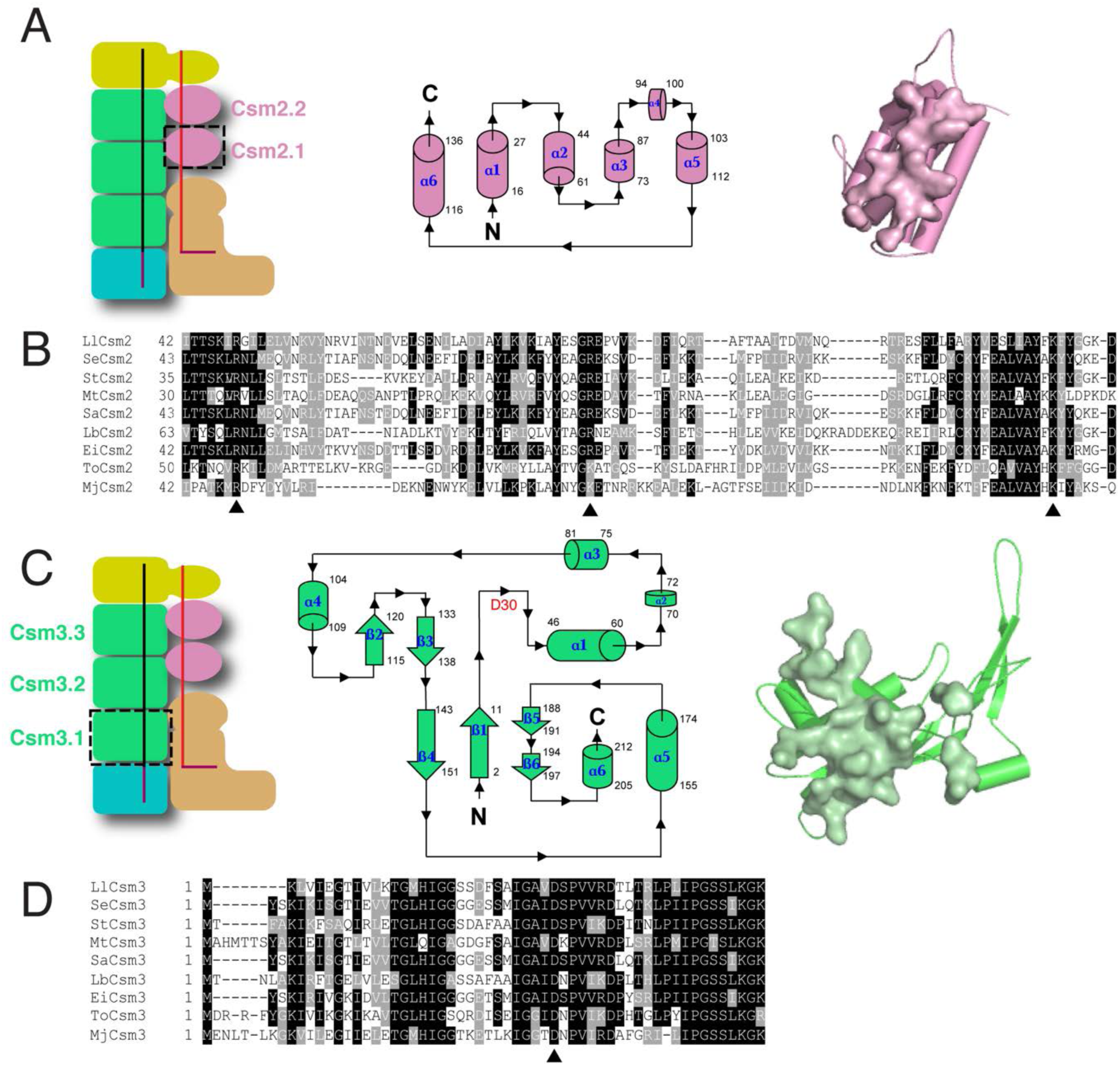
Topology, multiple sequence alignment and subunit interfaces of LlCsm2 and LlCsm3. Related to Figure 5. (A) Topology map of helical LlCsm2. Interface residues of Csm2 are identified to be 4 Å from its neighboring Csm2 subunit and are shown in surface representation. (B) Multiple sequence alignment of *Lactococcus lactis* Csm2 (LlCsm2) with bacterial Csm2 orthologs from *Staphylococcus epidermidis* (SeCsm2), *Streptococcus thermophilus* (StCsm2), *Mycobacterium tuberculosis* (MtCsm2), *Staphylococcus aureus* (SaCsm2), *Lactobacillus delbrueckii* subsp. *bulgaricus* (LbCsm2), *Enterococcus italicus* (EiCsm2) and archaeal orthologs from *Thermococcus onnurineus* (ToCsm2), *Methanocaldococcus jannaschii* (MjCsm2). Important residues are highlighted by black triangles. (C) Topology map of LlCsm3. The location of RNase catalytic site D30 is highlighted in red. Interface residues of Csm3 are identified to be 4 Å from its neighboring Csm3 subunit and are shown in surface representation. (D) Multiple sequence alignment of *Lactococcus lactis* Csm3 (LlCsm3) with bacterial Csm3 orthologs from *Staphylococcus epidermidis* (SeCsm3), *Streptococcus thermophilus* (StCsm3), *Mycobacterium tuberculosis* (MtCsm3), *Staphylococcus aureus* (SaCsm3), *Lactobacillus delbrueckii* subsp. *bulgaricus* (LbCsm3), *Enterococcus italicus* (EiCsm3) and archaeal orthologs from *Thermococcus onnurineus* (ToCsm3), *Methanocaldococcus jannaschii* (MjCsm3). The strongly conserved Aspartate residue important for target RNA cleavage is highlighted in black triangle.

**Figure S9.**
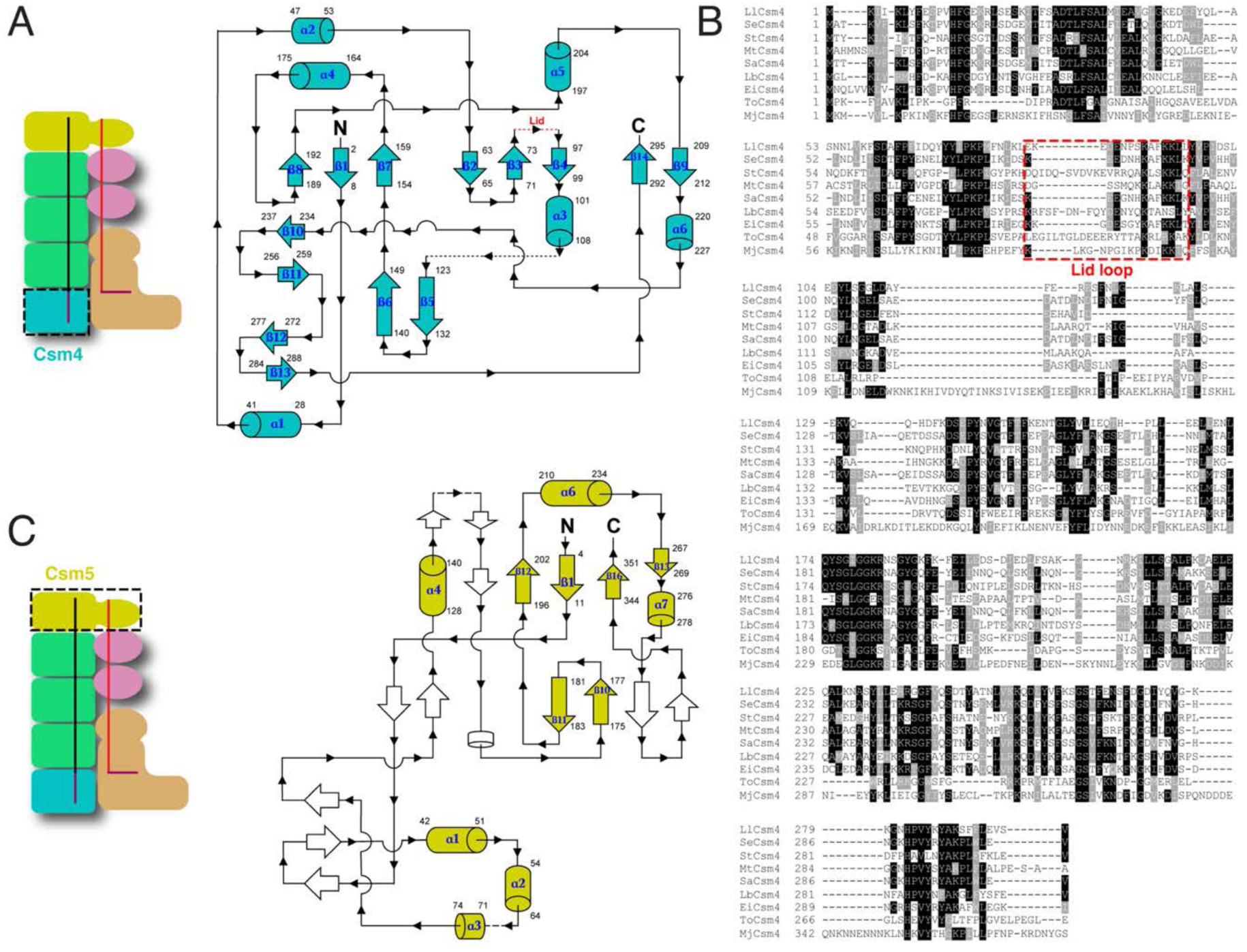
Topology, multiple sequence alignment of LlCsm4 and topology of LlCsm5. Related to Figures 2, 3 and 4. (A) Topology map of LlCsm4. The Lid loop is highlighted in red. (B) Multiple sequence alignment of *Lactococcus lactis* Csm4 (LlCsm4) with bacterial Csm4 orthologs from *Staphylococcus epidermidis* (SeCsm4), *Streptococcus thermophilus* (StCsm4), *Mycobacterium tuberculosis* (MtCsm4), *Staphylococcus aureus* (SaCsm4), *Lactobacillus delbrueckii* subsp. *bulgaricus* (LbCsm4), *Enterococcus italicus* (EiCsm4) and archaeal orthologs from *Thermococcus onnurineus* (ToCsm4), *Methanocaldococcus jannaschii* (MjCsm4). (C) Topology map of Csm5. Structural elements that could not be traced in the structure are left uncolored.

---

**Table S1:** Compilation of observations of plasmid interference assay and assessment of functionality of variants of LlCsm complex. Related to Figures 1 and S2. Wild type LlCsm harboring non-cognate crRNA (WT/NCR), wild type LlCsm harboring cognate crRNA (WT/CCR), LlCsm Csm3 Asp30Ala mutant (LlCsm (D30A)_3_), LlCsm Csm1 His13Ala Csm3 Asp30Ala double mutant (LlCsm (H13A)_1_(D30A)_3_), LlCsm Csm1 Asp14Asn Csm3 Asp30Ala double mutant (LlCsm (D14N)_1_(D30A)_3_), LlCsm Csm1 His13Ala Asp14Asn double mutant (LlCsm (H13A)_1_(D14N)_1_), LlCsm Csm1 (GGDD)_574-577_ to (GGAA)_574-577_ mutant (LlCsm (GGAA)_1_), Csm2-deleted LlCsm complex (ΔCsm2-LlCsm), Csm2-deleted LlCsm complex harboring Csm1 His13Ala Asp14Asn mutations (ΔCsm2-LlCsm (H13A)_1_(D14N)_1_) and Csm2-deleted LlCsm complex harboring Csm1 (GGDD)_574-577_ to (GGAA)_574-577_ mutation (ΔCsm2-LlCsm (GGAA)_1_) were tested.

| Llcsm variant | Plasmid Interference |
| --- | --- |
| WT/NCR | NO |
| WT/CCR | YES |
| Llcsm (D30A)<sub>3</sub> | YES |
| Llcsm (H13A)<sub>1</sub>(D30A)<sub>3</sub> | YES |
| Llcsm (D14N)<sub>1</sub>(D30A)<sub>3</sub> | YES |
| Llcsm (H13A)<sub>1</sub>(D14N)<sub>1</sub> | YES |
| Llcsm (GGAA)<sub>1</sub> | NO |
| ΔCsm2-Llcsm | NO |
| ΔCsm2-Llcsm (H13A)<sub>1</sub>(D14N)<sub>1</sub> | NO |
| ΔCsm2-Llcsm (GGAA)<sub>1</sub> | NO |

**Table S2:**
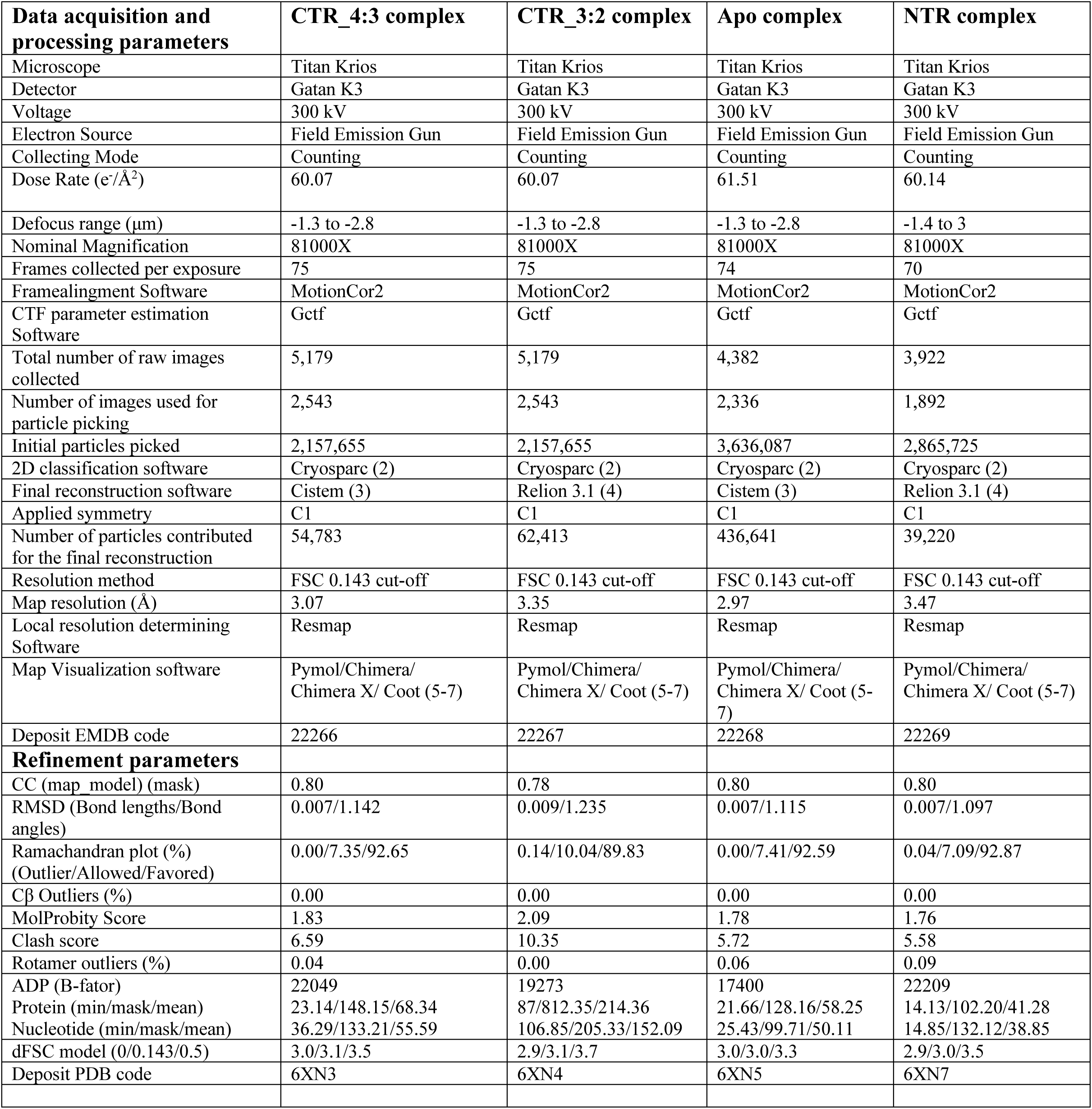
Statistics of cryo-EM data processing and model refinement. Related to Figures 2, 4 and 5, STAR Methods.

**Table S3:**
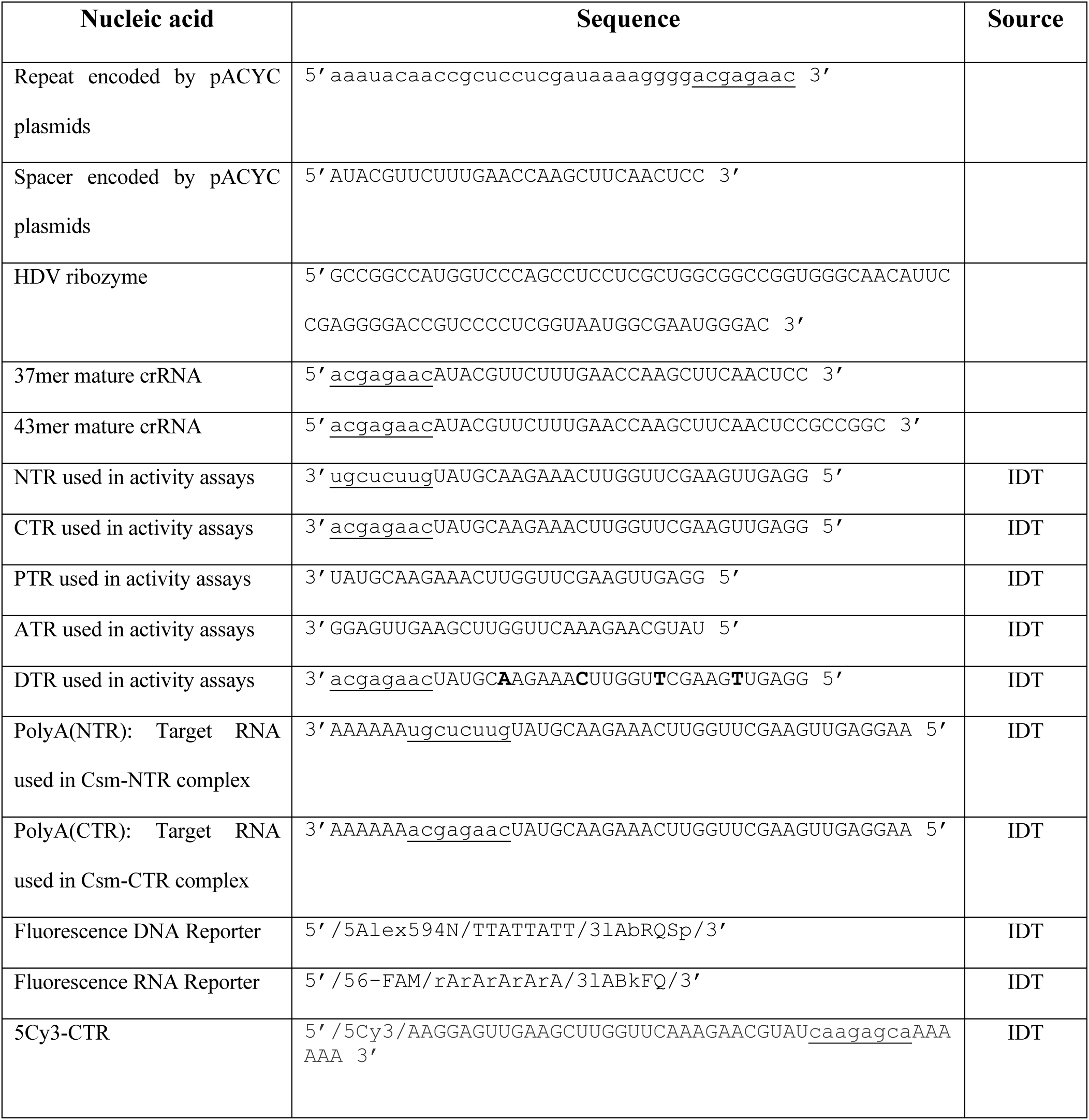
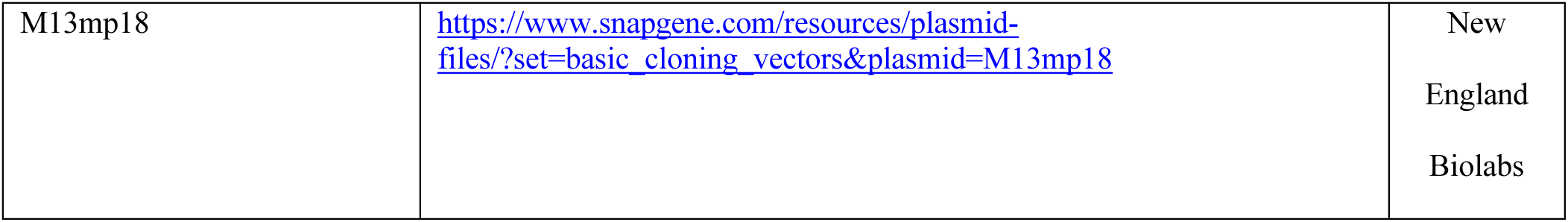
Nucleic acid sequences and oligonucleotides used in this study. Related to Figures 1, 4 and S1. Underlined regions indicate either the 5’-handle of the unprocessed/cas6-processed crRNA or the 3’-antitag of the target RNA. Bold in DTR (Deoxyribonucleotide target RNA) highlights Csm3-mediated ribonucleotide cleavage sites replaced by deoxyribonucleotides.

**Table S4:**
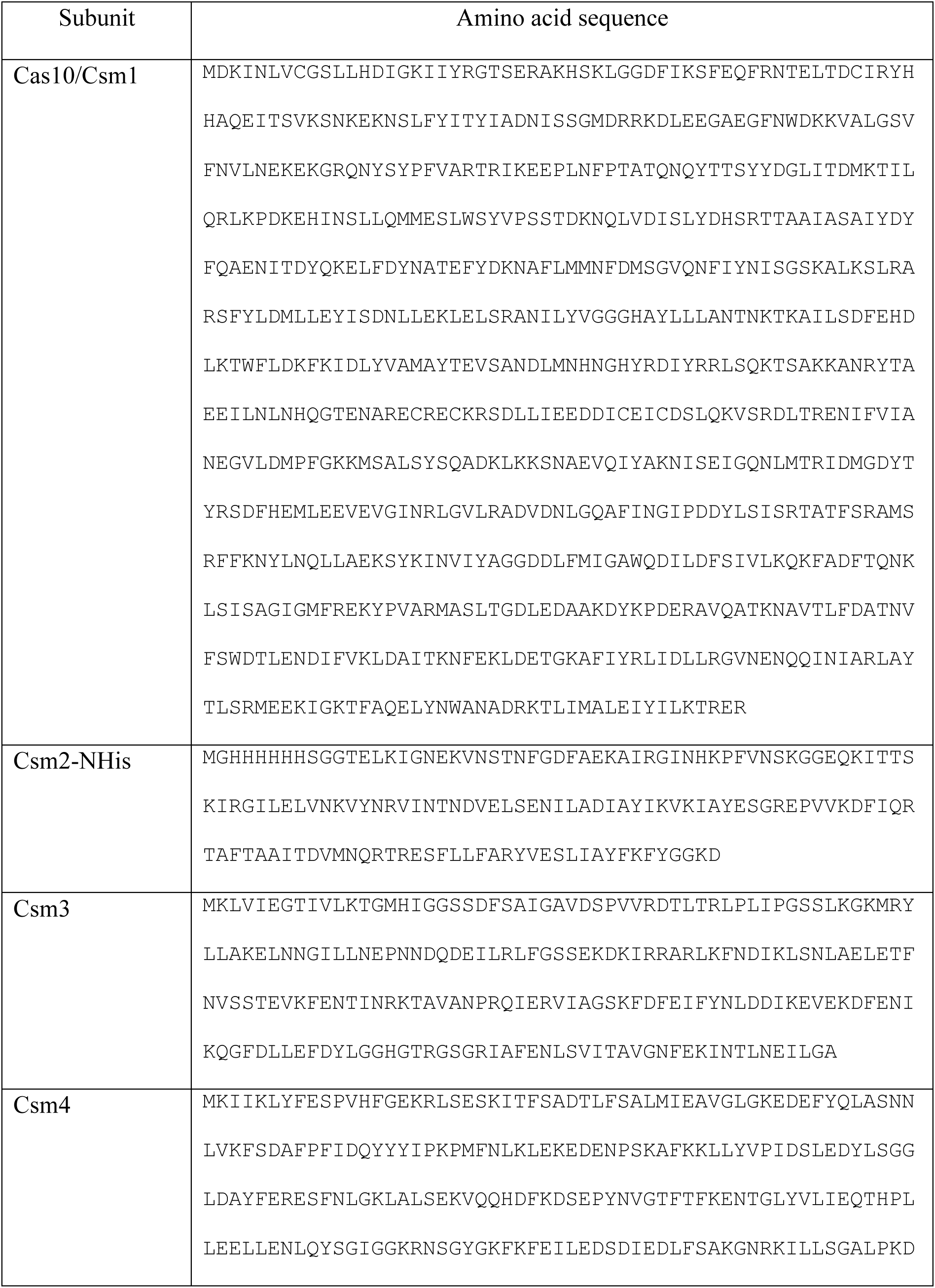
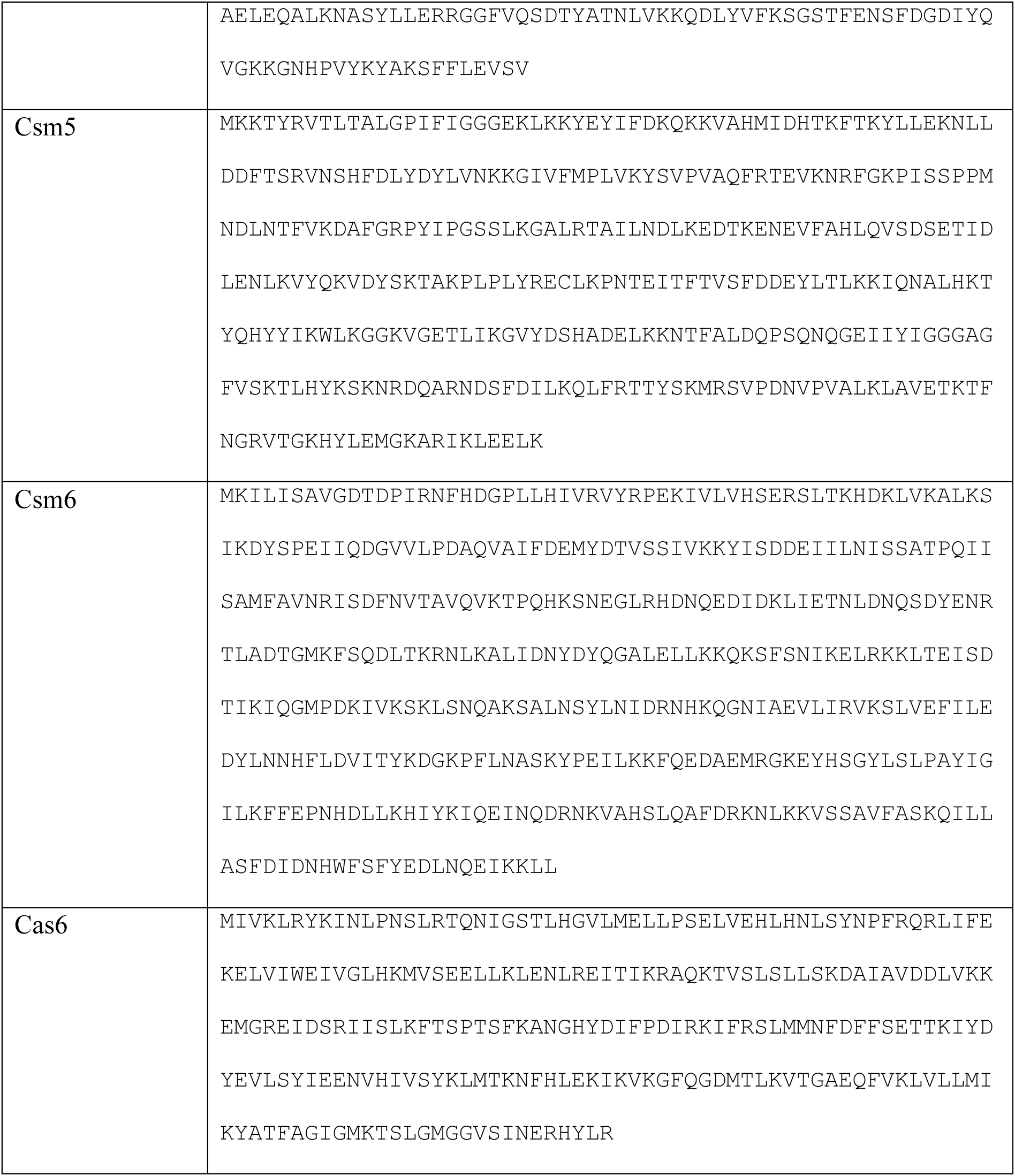
Amino acid sequences of *Lactococcus lactis* CRISPR-Cas (Csm) system. Related to Figures 1, 4 and S1.

